# Genomic architecture of 5S rDNA cluster and its variations within and between species

**DOI:** 10.1101/2021.02.17.431734

**Authors:** Qiutao Ding, Runsheng Li, Xiaoliang Ren, Lu-yan Chan, Vincy W. S. Ho, Dongying Xie, Pohao Ye, Zhongying Zhao

**Affiliations:** Department of Biology, Hong Kong Baptist University, Hong Kong; State Key Laboratory of Environmental and Biological Analysis, Hong Kong Baptist University, Hong Kong, China SAR

**Keywords:** *Caenorhabditis elegans*, *C. briggsae*, rDNA cluster, 5S, 18S-5.8S-26S, Nanopore sequencing

## Abstract

Ribosomal genes (rDNAs) are arranged in purely tandem repeats, preventing them from being reliably assembled onto chromosome. The uncertainty of rDNA genomic structure presents a significant barrier for studying their function and evolution. Here, we generate ultra-long Nanopore and short NGS reads to delineate the architecture and variation of the 5S rDNA cluster in the different strains of *C. elegans* and *C. briggsae*. We classify the individual rDNA units into 25 types based on the unique sequence variations in each unit of *C. elegans* (N2). We next perform manual assembly of the cluster using the long reads that carry these units, which led to an assembly of rDNA cluster consisting of up to 167 5S rDNA units. The ordering and copy number of various rDNA units are indicative of separation time between strains. Surprisingly, we observed a drastically lower level of variation in the 5S rDNA cluster in the *C. elegans* CB4856 and *C. briggsae* AF16 strains than *C. elegans* N2 strain, suggesting a unique mechanism in maintaining the rDNA cluster stability in the N2. Single-copy transgenes landed into the rDNA cluster shows the expected expression in the soma, supporting that rDNA genomic environment is transcriptionally compatible with RNA polymerase II. Delineating the structure and variation of rDNA cluster paves the way for its functional and evolutionary studies.

## Introduction

Ribosomal RNAs (rRNAs) as the components of ribosome play a critical role in protein synthesis. Eukaryotic rRNAs are encoded by ribosomal DNAs (rDNAs) that are arranged in tandem within rDNA cluster. There are four rRNA genes, i.e., 5S rRNA, 18S rRNA, 5.8S rRNA, and 28S rRNA. The 5S rDNA cluster is usually arranged as tandem repeats that are away from the remaining three genes in most species with a few exceptions, including yeast [1]. The 18S, 5.8S, and 28S rRNAs are produced as a single transcript using 45S rDNA as a template, which is also arranged as tandem array in the genome. The transcript is processed into three individual ones following transcription [2]. In contrast to most mRNAs and microRNAs that are produced with RNA polymerase II (Pol II) [3], the 45S rRNAs are transcribed by RNA polymerase I (Pol I), and the 5S rRNAs are transcribed by RNA polymerase III (Pol III) along with tRNA. Intriguingly, the rRNAs made with RNA pol II was able to rescue the phenotype of an rDNA deletion mutant in yeast [4], indicating that rRNAs transcribed by RNA pol II are functional. However, it remains unclear whether the genomic environment of rDNAs consisting of tandem repeats is permissive for mRNA transcription.

The rDNA copy number is known to be variable between cells, or individuals with different age [5–7]. The copy number variation (CNV) was found to be coupled with carcinogenesis [6]. Interestingly, the extrachromosomal rDNA circles derived from chromosomal rDNA repeats can be reintroduced into the host genome in a dosage-dependent behavior [8], further complicating the copy number variation. The rDNA CNV between different wild isolates or mutated strains of *C. elegans* has been estimated with <u>n</u>ext-<u>g</u>eneration <u>s</u>equencing (NGS) reads and quantitative PCR [5, 9], ranging from 33 to 245 copies for the 45S rDNA and 39 to 438 copies for the 5S rDNA. However, the rDNA CNV during development has not been reported in *C. elegans* or in other nematodes. In addition to CNV, sequence variation is also noted in the rDNAs of the same species [10, 11].

The sequences of rDNA and its <u>n</u>on-transcribed <u>s</u>equence (NTS) are found to have polymorphisms in eukaryotic species, including, <u>s</u>ingle-<u>n</u>ucleotide <u>p</u>olymorphisms (SNPs) and small <u>in</u>sertion or <u>del</u>etion (INDEL). For example, in the mouse and human, the INDELs ranging from 1- to 12 bps in rDNA were frequently identified between chromosomes, tissues, individuals, and families [12, 13]. Similar polymorphisms in rDNA were also identified in yeast [14], fly [15] and plants [16]. Unexpectedly, *C. elegans* carries only one type of 5S rDNA unit with few SNPs in its coding sequence [10, 17], whereas its related species, *C. briggsae*, carries two distinct types of 5S rDNA units with distinctive orientation of 5S rDNA relative to <u>s</u>plicing leader 1 (SL1) [18, 19]. Whether there are any 5S rDNA variants in the NTS region of nematode species has not been thoroughly investigated.

NGS techniques have been intensively used to assemble genome across species in the past two decades, leading to an exponential increase of genomic data across species. However, the genome assembly produced with NGS only is usually fragmented due to the presence of repetitive sequences, especially in those regions consisting of highly tandem repeats such as centromeres and rDNAs. Therefore, these tandem repeats are commonly included in various contigs that are unable to be assigned onto precise location of chromosome. The repetitive sequences create a huge challenge for genome assembly using NGS reads because of their relatively short read length ranging from 100 to 200 bps. Therefore, extra efforts have been made to improve the continuity of an assembly, including mate-pair sequencing of the ends from a large genomic fragment [20], incorporation of genetic markers [21] or chromatin configuration (Hi-C) [22], or using the long reads synthesized with the NGS short reads [23]. These steps have significantly improved the continuity of genome assembly, especially for those relatively small genomes. *C. elegans’* isogenic genome is the first metazoan genome that was assembled using Sanger sequencing reads coupled with physical mapping, leading to an exceptionally high contiguity [21]. It barely contains any gaps except in the rDNA clusters and telomere sequences. However, the high mapping costs prevent its universal application to other species. The genome assembly of its companion species, *C. briggsae*, was generated using shotgun sequencing coupled with scaffolding with end sequencing of <u>b</u>acterial <u>a</u>rtificial <u>c</u>hromosome (BAC) and fosmids [19]. The resulting contigs or supercontigs were assembled onto chromosomes using genetic markers [24] or synthetic long reads (SLR) coupled with Hi-C [23]. However, these efforts failed to resolve the localization and genomic organization of rDNA clusters. Delineation of the genomic architecture and localization of rDNA clusters is needed for studying the evolution, function, and regulation of ribosomal genes [25–28].

Third-<u>g</u>eneration sequencing (TGS) techniques, including Oxford Nanopore Technologies (ONT) Nanopore sequencing and PacBio Single Molecule, Real-Time (SMRT) sequencing, overcome the intrinsic limitation of the short-read by generating ultra-long reads with limited sequencing bias [29], which is expected to facilitate genome assembly with an improved continuity by the inclusion of more repetitive sequences [30–32]. Importantly, the amplification-free TGS enables researchers to directly sequence DNA or RNA with a reduced sequence bias [33]. Due to its ultra-long length, TGS has recently been used to re-sequence *C. elegans* genome, which recovered substantially more repetitive sequences and revealed chromosomal rearrangements and structural variations between strains [30–32]. However, these assemblies were not able to resolve the genomic structure of 5S rDNA and 45S rDNA cluster.

Here, we characterized the genomic architecture of the 5S rDNA cluster in both *C. elegans* and *C. briggsae* using both ONT sequencing and NGS reads. Aided by the reads, we identified various reproducible sequence variations in the 5S rDNA unit in both species, which allowed us to generate an assembly of 5S rDNA cluster carrying up to at least 167 repetitive units. The ONT reads also permitted the determination of genomic localization of rDNAs in *C. briggsae*. We observed strain-specific composition and CNV of the 5S rDNA units that are indicative of separation time among *C. elegans* strains. Our functional characterization of the rDNA cluster indicates the genomic environment of rDNA cluster is transcriptionally compatible for RNA polymerase II at least in the somatic tissues. Our structural and functional characterization of the rDNA clusters lays a foundation for further characterization of the rDNA function, regulation and evolution.

## Results

### Genomic architecture of the 5S rDNA cluster

To gain an initial idea of the genomic architecture of rDNA cluster, starting from the existing *C. elegans* N2 genome assembly WBcel235 [21], we focused on the 5S rDNA cluster on the chromosome V for manual finishing due to its relatively short size and the well-characterized boundary sequences (Fig. S1). We generated ∼1.8 million ONT DNA reads with an N50 from 18 to 31 Kbp from three developmental stages of *C. elegans* N2, i.e. embryo (EMB), L1 larvae (L1), and young adult (YA) stages (Table 1), which were mapped against the reference genome WBcel235 [34]. As expected, the mapping results showed a drastic increase in read coverage of 5S rDNA compared with its flanking sequences (Fig. S2a). The flanking sequences of the 5S rDNA cluster were identified using the ONT reads that spanned the rDNA repeats and the unique sequences on both sides of the cluster (Fig. 1c). Given that the genomic structure of rDNA cluster has not been resolved in any species due to its extremely repetitive characteristics, we set out to investigate whether there are any sequence variants in the rDNA units that could be harnessed to assemble the entire cluster by sequencing of three developmental stages of *C. elegans* (Table 2). Unexpectedly, we not only confirmed the presence of the canonical 5S rDNA unit (referred to as unit 1.1 hereafter) in *C. elegans*, but also identified numerous novel variants of the 5S rDNA unit that are reproducibly arranged relative to one another in the ONT reads. We chose a subset of the variants of 5S rDNA unit to facilitate our assembly of 5S rDNA cluster (Fig. 1a, b, and Table 2). We classified the rDNA units into three major categories, i.e., 1-3, by the presence or absence of two unique deletions, i.e., 99_102del, 780_809del, representing a deletion of 4 and 30 bps, respectively, from the indicated positions relative to the canonical 5S rDNA unit 1.1. Each category of unit was further classified into different types based on other sequence variations, mostly SNPs, on top of the two deletions established with the existing NGS reads (Table 2). The relative proportion of each 5S rDNA variant with unique SNP/INDEL was confirmed with the NGS reads [35] (Fig. 1b).

**Fig. 1.**
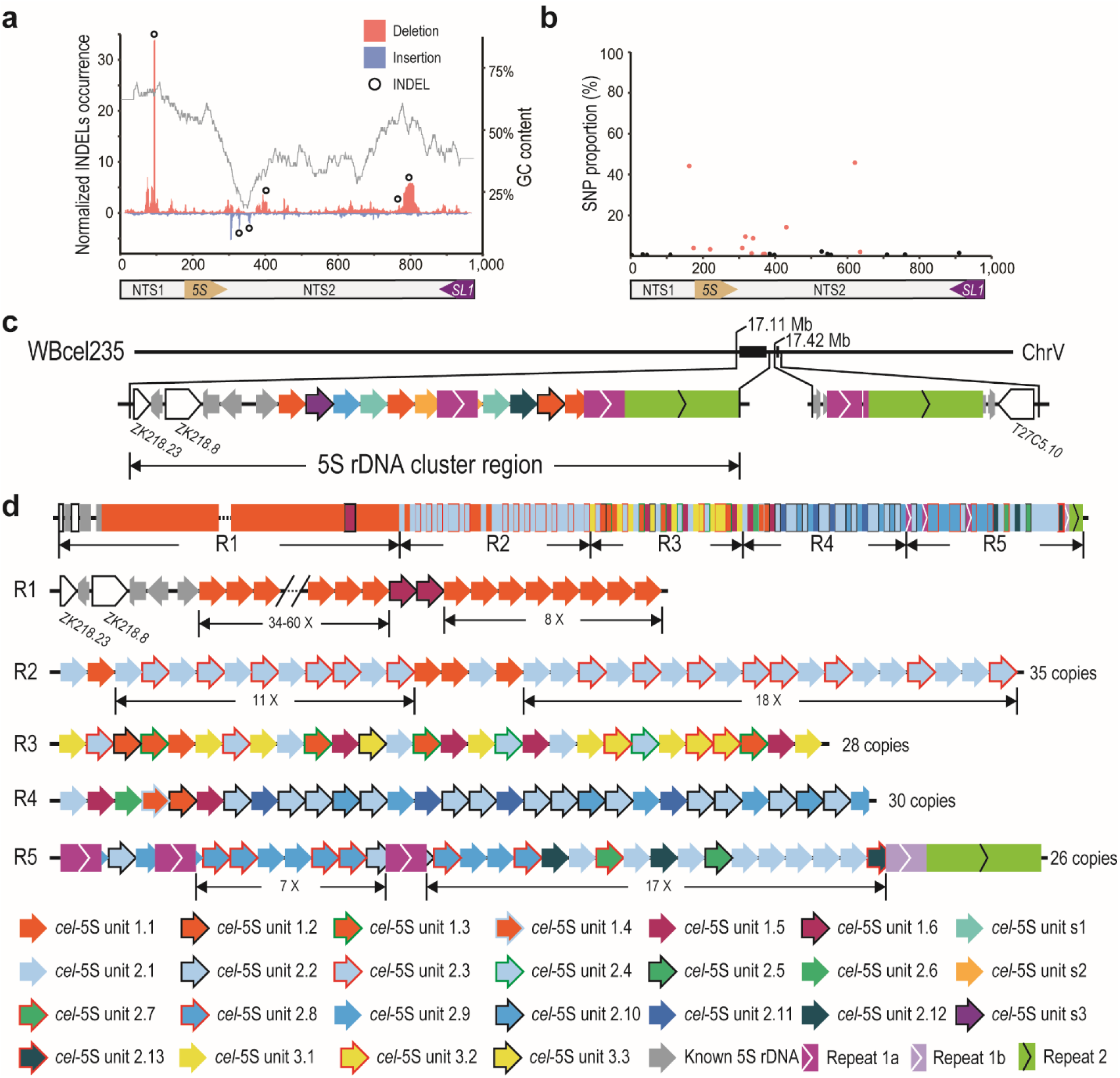
Structure of the *C. elegans* (N2) 5S rDNA cluster. (a) INDELs identified with Nanopore reads within 5S rDNA unit. Shown are normalized INDEL occurrences along with GC content. Deletion and insertion identified with Nanopore raw reads are shown in red and blue, respectively. Cross-validated INDELs used in the subsequent analysis are indicated with black circles (see Methods). (b) SNPs in the 5S rDNA are identified with existing NGS data. SNPs present or absent in new rDNA variants are colored in red and black, respectively. (c) Structure of the 5S rDNA-containing regions on the chromosome V in the current *C. elegans* N2 reference genome (WBcel235). (d) Structure of the 5S rDNA cluster assembled by ONT reads, which carries a total of at least 167 partial or complete units. The cluster is divided into 5 regions (R1-5) based on the SNPs and INDELs present in each unit or the position relative to Repeat 1a. Newly identified rDNA units or unique repeats are differentially color coded (see Table 2). Variation in rDNA copy number is indicated with dash line. Note that three Repeat 1a are inserted into 5S rDNA unit at same position within R5.

**Table 1.** Nead statistics.

| **Library name** | **Total**<br>**number of**<br>**reads** | **Total**<br>**bases**<br>**(Gbp)** | **Mean**<br>**length**<br>**(bp)** | **Median**<br>**length**<br>**(bp)** | **N50**<br>**length**<br>**(bp)** | **Max**<br>**mapped**<br>**length (bp)** |
| --- | --- | --- | --- | --- | --- | --- |
| **N2-EMB** | 789,871 | 10.8 | 13,724 | 13,084 | 18,558 | 163,153 |
| **N2-L1** | 199,712 | 3.7 | 18,300 | 13,605 | 31,265 | 196,902 |
| **N2-YA** | 822,902 | 9.3 | 11,341 | 9,187 | 19,566 | 174,664 |
| **AF16-YA** | 1,433,280 | 11.1 | 7,724 | 4,148 | 15,427 | 182,506 |
| **ZZY0600** | 870,874 | 12.6 | 14,479 | 10,248 | 25,074 | 247,180 |
| **ZZY0603** | 2,696,939 | 12.9 | 4,785 | 2,463 | 9,429 | 252,751 |
| **ZZY0653** | 60,187 | 0.6 | 10,720 | 2,409 | 27,723 | 139,839 |
| **CB4856** | 2,294,403 | 15.1 | 6,562 | 2,347 | 19,197 | 382,430 |

EMB: Mix-staged embryos; L1: Larval stage 1; YA: Young adult

**Table 2.** List of the variants of 5S rDNA unit in *C. elegans* N2 used in this study.

| Variant | Size (bp) | Copy number | Sequence variation relative to *cel*-5S unit 1.1 |
| --- | --- | --- | --- |
| unit 1.1 | 976 | Dynamic | Not applicable |
| unit 1.2 | 976 | 2 | 621T>G |
| unit 1.3 | 976 | 4 | 220C>A, 621T>G |
| unit 1.4 | 976 | 1 | 162C>G, 621T>G |
| unit 1.5 | 980 | 6 | 309T>C, 318T>C, 325_326insCAAT, 329G>T, 332T>G, 621T>G |
| unit 1.6 | 971 | 2 | 766_771delinsC |
| unit 2.1 | 972 | 29 | 99_102del, 162C>G, 621T>G |
| unit 2.2 | 972 | 15 | 99_102del, 162C>G, 431T>G, 621T>G |
| unit 2.3 | 972 | 16 | 99_102del, 162C>G |
| unit 2.4 | 972 | 2 | 99_102del, 162C>G, 220C>A, 621T>G |
| unit 2.5 | 976 | 1 | 99_102del, 162C>G, 335G>C, 407C>T, 621T>G |
| unit 2.6 | 976 | 1 | 99_102del, 162C>G, 309T>C, 318T>C, 325_326insCAAT, 329G>T, 332T>G, 621T>G |
| unit 2.7 | 976 | 1 | 99_102del, 162C>G, 318T>C, 325_326insCAAT, 329G>A, 332T>G, 339C>T, 621T>G |
| unit 2.8 | 976 | 6 | 99_102del, 162C>G, 318T>C, 325_326insCAAT, 329G>A, 332T>G, 339C>T, 431T>G, 621T>G, 636T>G |
| unit 2.9 | 976 | 9 | 99_102del, 162C>G, 318T>C, 325_326insCAAT, 329G>T, 332T>G, 339C>T, 621T>G |
| unit 2.10 | 976 | 3 | 99_102del, 162C>G, 318T>C, 325_326insCAAT, 329G>T, 332T>G, 339C>T, 431T>G, 621T>G |
| unit 2.11 | 963 | 4 | 99_102del, 162C>G, 318T>C, 325_326insCAAT, 329G>T, 332T>G, 339C>T, 393_405del, 431T>G, 621T>G |
| unit 2.12 | 982 | 3 | 99_102del, 162C>G, 309T>C, 318T>C, 325_326insCAAT, 329G>T, 332T>G, 354_355insGGTATT, 367A>T, 371T>A, 621T>G |
| unit 2.13 | 984 | 1 | 99_102del, 162C>G, 309T>C, 318T>C, 325_326insCAAT, 329G>T, 332T>G, 354_355insGGTATT, 367A>T, 371T>A, 621T>G, 718_719insGA |
| unit 3.1 | 942 | 7 | 780_809del, 99_102del, 162C>G, 621T>G |
| unit 3.2 | 946 | 3 | 780_809del, 621T>G |
| unit 3.3 | 950 | 1 | 780_809del, 220C>A, 309T>C, 318T>C, 325_326insCAAT, 329G>T, 332T>G, 621T>G |
| unit s1* | 975 | 0 | 99_102del, 162C>G, 325_326insA, 393delC, 621T>G |
| unit s2* | 981 | 0 | 354_355insGGTATT, 367A>T, 371T>A, 545G>A, 621T>G |
| unit s3* | 972 | 0 | 325_326insCAAT, 329G>A, 332T>G, 339C>T, 431T>G, 621T>G |

\*Combinations of variants in s1-s3 are not identified in the 5S rDNA cluster. Del: deletion. Ins: insertion. Delins: deletion followed by insertion.

The genomic organization of rDNA units was resolved through tiling of ONT reads from both orientations by taking advantage of different combinations of rDNA unit variants and other types of repeats present in the rDNA cluster (Fig. 1a-c, Fig. S1, Table 2, and Table S1). Consequently, we were able to generate a contig that carries a total of at least 167 copies of 5S rDNA units (Fig. 1d), including at least 47 copies of canonical rDNA unit (unit 1.1), 15, 90 and 11 copies of unit 1, 2 and 3 variants, respectively, and 4 copies of existing 5S rDNA unit. In addition, there are 3 copies of existing non-rDNA repeat (referred to as Repeat 1a, 1b, and 2) (Table S2) in the cluster. The rDNA cluster were divided into five regions (R1-5) based on the number and composition of the 5S rDNA units. The results show that 5S rDNA cluster consists of various unit variants arranged in reproducible order in our N2 strain. Availability of the detailed structure of 5S rDNA cluster is expected to facilitate functional and evolutionary analysis of rDNAs.

### Structural variations of 5S rDNA cluster between our N2 and its derived *C. elegans* strains

Given that the relatively stable number and genomic organization of 5S rDNA variants in the ONT reads derived from our *C. elegans* N2, we wondered to what extent the arrangement and copy number of the rDNA units are conserved between our N2 and other N2-derived strains that had been separated from one another for different times. To this end, we generated ∼0.9 and ∼2.7 million ONT reads for two transgenic strains (ZZY0600 and ZZY0603), each carrying a single copy of transgene associated with 5S rDNA sequences (Fig. 2a, b) generated using *miniMos* technique [36] in the background of *unc-19* mutant allele *tm4063* [37]. Reads with transgene sequences helped us elongate the assembly of 5S rDNA cluster but still failed to span the entire cluster region. Three rDNA structural variations were found between the 5S rDNA clusters of our N2 and two transgenic strains (Fig. 2c-e). In addition to our sequenced data from N2 and the transgenic strains, we also used existing ONT reads generated from other N2-derived strains [30, 31] to further evaluate the variation in the 5S rDNA clusters because the two N2 strains were separated from each other and individually maintained for at least 10 years. Intriguingly, we observed variations across Region 1-4. Extent of variation is indicative of separation time between each other, i.e., the longer time the two strains are separated between each other, the more variations are found between the structures of their 5S rDNA clusters. For example, a fragment consisting of one copy of unit 3 and two copies of unit 1 is missing in the Region 3 of our N2 relative to all the remaining strains (Fig. 2d). More variations in the copy number of *C. elegans* 5S unit (*cel*-5S unit) 1 were observed in the Region 4 (Fig. 2e). Our N2 contains 30 copies of rDNA unit, whereas the two transgenic strains derived from the same starting strain have 32 copies, and the strain VC2010 and its recent derivative PD1074 both carry 35 copies. However, the VC2010 [31] gains an extra four copies of unit 2 after its separation from its derived strain PD1074 (Fig. 2f) [30]. This apparent association of rDNA type and/or exact copy number with separation time raises the possibility of using the variation in barcoding the strains that are freshly separated from one another (Fig. 2g).

**Fig. 2.**
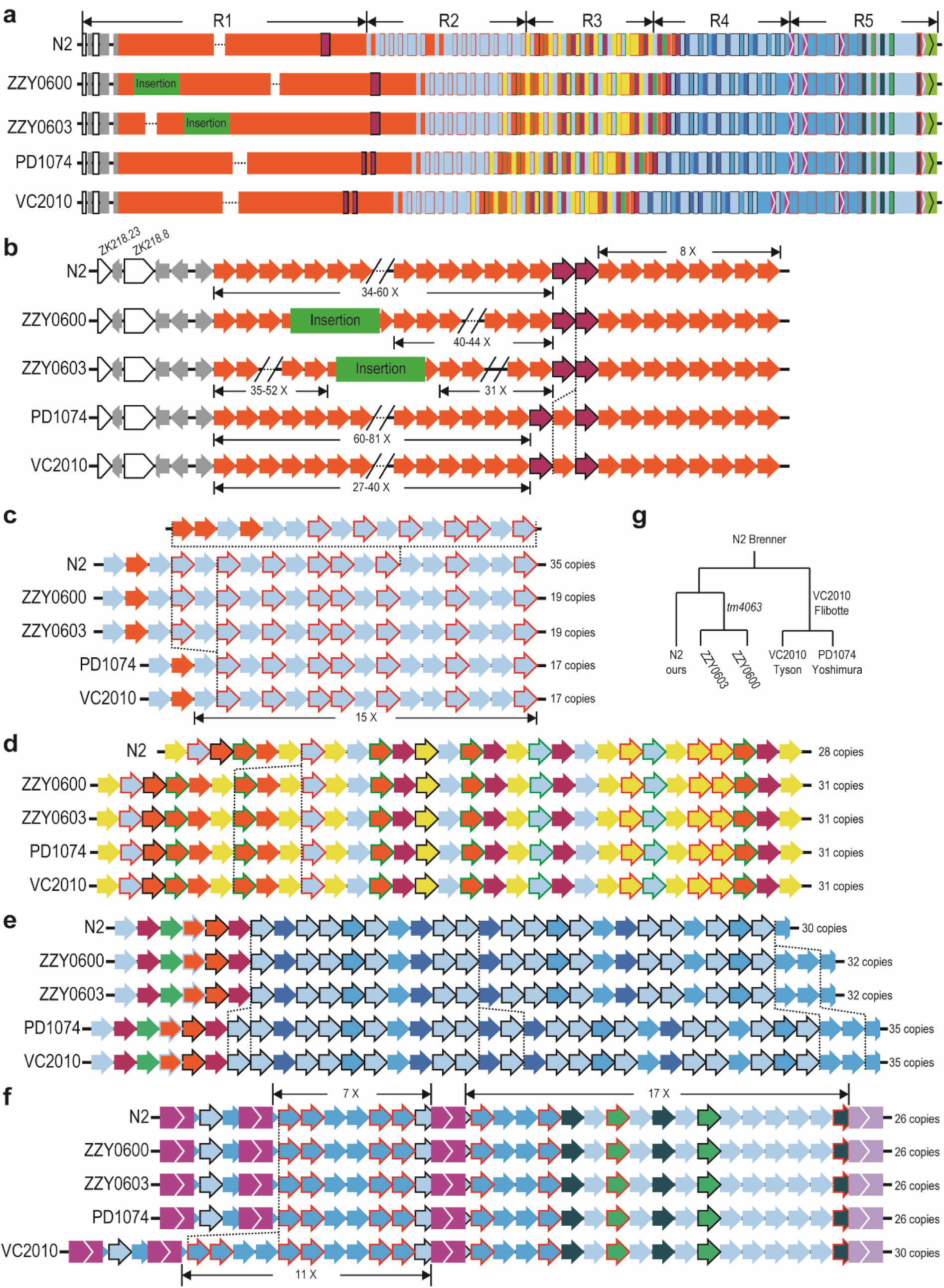
Structural variations within 5S rDNA cluster between our *C. elegans* N2 and other N2-derived strains. (a) Overview of the structures of 5S rDNA clusters for five strains as shown in Fig. 1d. Strain names are indicated on the left. Position and size of transgenic insertions are indicated in scale. (b) Comparison of unit compositions and estimated copy number in R1. Identified variation in unit composition is highlighted with vertical dashed line. (c-f) Comparison of unit compositions in R2 (c), R3 (d), R4 (e), and R5 (f) as in (b). (g) Ancestry of the strains based on strain history. Our N2 was shipped from Waterston lab in 2010. PD1074 was a recent derivative of VC2010 that was derived from a separate N2 in Don Moerman lab. ZZY0600 and ZZY0603 were generated by transgene insertion into *unc-119*(*tm4063*) worms, which was derived from another *C. elegans* N2 in Mitani lab.

### Largely uniform composition of 5S rDNA unit in the 5S rDNA cluster of *C. elegans* Hawaii strain and *C. briggsae* wild isolate AF16

To further examine the structural variations in rDNA cluster between N2 and more distantly related *C. elegans* strains, we focused on the comparison between N2 and CB4856, a Hawaiian strain of *C. elegans* that is one of the most divergent from the strain N2 [38]. To this end, we generated ∼2.3 million ONT reads using CB4856 animals (Table 1), which were used to assemble the 5S rDNA cluster of CB4856 in a way similar to that used for the N2 (Fig. 3). Surprisingly, we found that the canonical *C. elegans* 5S rDNA unit, i.e., *cel*-5S unit 1.1, one of the most predominant forms in N2, and *cel*-5S unit 3 were absent in the CB4856 genome using a combination of existing NGS reads with our ONT reads for CB4856 (Fig. 3a-e, Table 1, and Table S3). Remarkably, the occurrences of SNP and INDEL identified in 5S rDNA unit are much lower in the CB4856 than in the N2 strain (Fig. 3a-b). All the units in CB4856 belong to the category 2 due to the presence of a 4 bp-deletion (Fig. 3a and Table 2). They can be further divided into six subtypes versus the 13 in the N2 (Table 2 and Table S3). Only two subtypes out of the six, i.e., unit 2.1 and 2.6, are shared between the two strains. Notably, the entire 5S rDNA cluster is dominated by two rDNA subtypes, i.e., units 2.14 and 2.15, with the former as the predominant member (Fig. 3c-d). The presence of relatively uniform rDNA units in CB4856 is in sharp contrast to the mosaic compositions of rDNA units in the N2 (Fig. 3c). The *cel*-5S unit variants 2.14 and 2.17 were interrupted by the Repeat 1a and 2 at the identical unit position (947-953 bp) in CB4856 (Fig. 3d), raising the possibility of their common origin. Given the presence of *cel*-5S unit 3 in N2 but not in CB4856 that carries a 30 bp deletion (Fig. 1-3, Table 2, and Table S3), we evaluated the distribution dynamics of the deletion using the existing NGS data from 330 *C. elegans* wild-isolates [35]. The result confirmed the presence of the unit 3 in N2 and other 163 its related strains but not in the remaining strains, including CB4856 (Fig. 3e, Fig. S3, Table S4). It also showed that this unique deletion had undergone multiple times of gain or loss between strains, suggesting a high turnover rate of 5S rDNA variation.

**Fig. 3.**
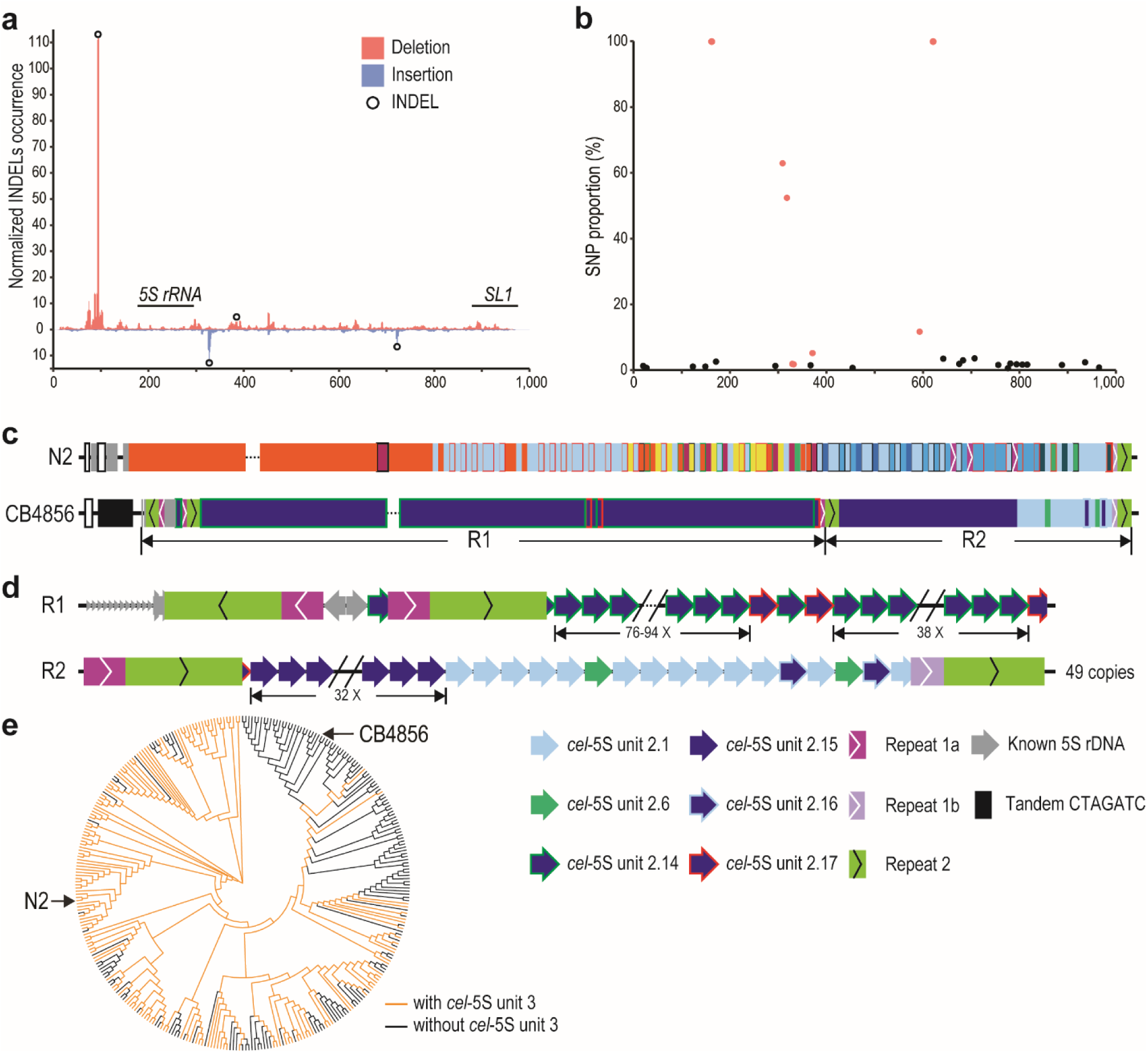
Structural variations within 5S rDNA clusters between *C. elegans* N2 and CB4856 strains. (a) INDELs identified with CB4856 Nanopore reads within 5S rDNA unit as in Fig. 1a. Cross-validated INDELs used in the subsequent analysis are indicated with black circles (see Methods). (b) SNPs in the 5S rDNA are identified with existing NGS data as in Fig. 1b. SNPs present or absent in new rDNA variants are colored in red and black, respectively. (c) Overview of the structures of 5S rDNA clusters between N2 and CB4856 as shown in Fig. 1d. Note the differences between the two, including lack of unit 1.1 (red) predominately seen in N2, whereas the unit (*cel*-5S 2.14) is unique to and predominately seen in CB4856. (d) Structure of the *C. elegans* CB4856 5S rDNA cluster. The Repeat 1a and 1b are shown as in Fig. 1d. (e) Evolutionary trajectory of the 5S rDNA unit 3 in *C. elegans*. Shown is a phylogenetic tree generated using SNPs from 330 *C. elegans* wild isolates. Strains with or without *cel*-5S unit 3 are colored in yellow and black, respectively. For simplicity, only N2 and CB4856 are shown (see Fig. S3 for a full list of strain names).

To further examine to what extent the structure of the 5S rDNA cluster is conserved between species, we generated ∼1.4 million of ONT reads (approximately 91× coverage) from *C. briggsae* AF16 young adults with an N50 of ∼15.4 kb and ∼39 million of paired end NGS reads of 150 bps in length from mix-staged *C. briggsae* animals. To locate the flanking sequences of 5S rDNA cluster in the *C. briggsae* genome, we combined the ONT reads with the previous SLR reads [23] to generate an AF16 genome assembly using Miniasm [39], followed by polishing with Racon [40]. After removal of bacterial genome and duplicated contigs, this draft assembly contains 20 contigs with summed size of approximately104 Mbp (Fig. 4a). The contigs were ordered and oriented relative to one another with the reference to CB4 [24] (Fig. 4b). Evaluation using BUSCO [41] revealed the completeness of this genome assembly was comparable to that of the *C. elegans* N2 genome (Fig. 4c). The *C. briggsae* genome was known to contain two divergent 5S rDNA units with an opposite orientation of SL1 relative to 5S rDNA coding sequence. They were referred to as *cbr*-5S unit 1.1 and 2.1, respectively (Fig. 5a-c), which were previously placed onto two separate locations on chromosome [28] (Fig. 5d). With two SNPs in *C. briggsae* 5S unit (*cbr*-5S unit) 1.1 (195G>T and 674G>T) and one deletion identified in the NGS data relative to *cbr*-5S unit 1.1 and 2.1 (382_440del), respectively (Table S5), we classified the *C. briggsae* 5S units into six types, i.e., unit 1.1-1.4 and unit 2.1-2.2, and generated the 5S rDNA cluster assembly in *C. briggsae* (AF16) in a way similar to that we did in *C. elegans*. Our new genome assembly and Hi-C data [28] supported that all the six divergent 5S rDNA units were located within a single location in the *C. briggsae* genome (Fig. 5e and Fig. S4b). The results also showed that *C. briggsae* 5S rDNA cluster mainly consisted of four type of units, i.e., 1.1, 1.2 1.4 and 2.1 (Fig. 5e). In summary, although the variations in sequence and copy number of 5S rDNA unit are quite common in *C. elegans* N2 and its derived strains, the 5S rDNA unit is largely uniform in *C. elegans* Hawaii strain (CB4856) and *C. briggsae* wild isolate (AF16), suggesting that the N2 is unique in maintaining the stability of its 5S rDNA cluster.

**Fig. 4.**
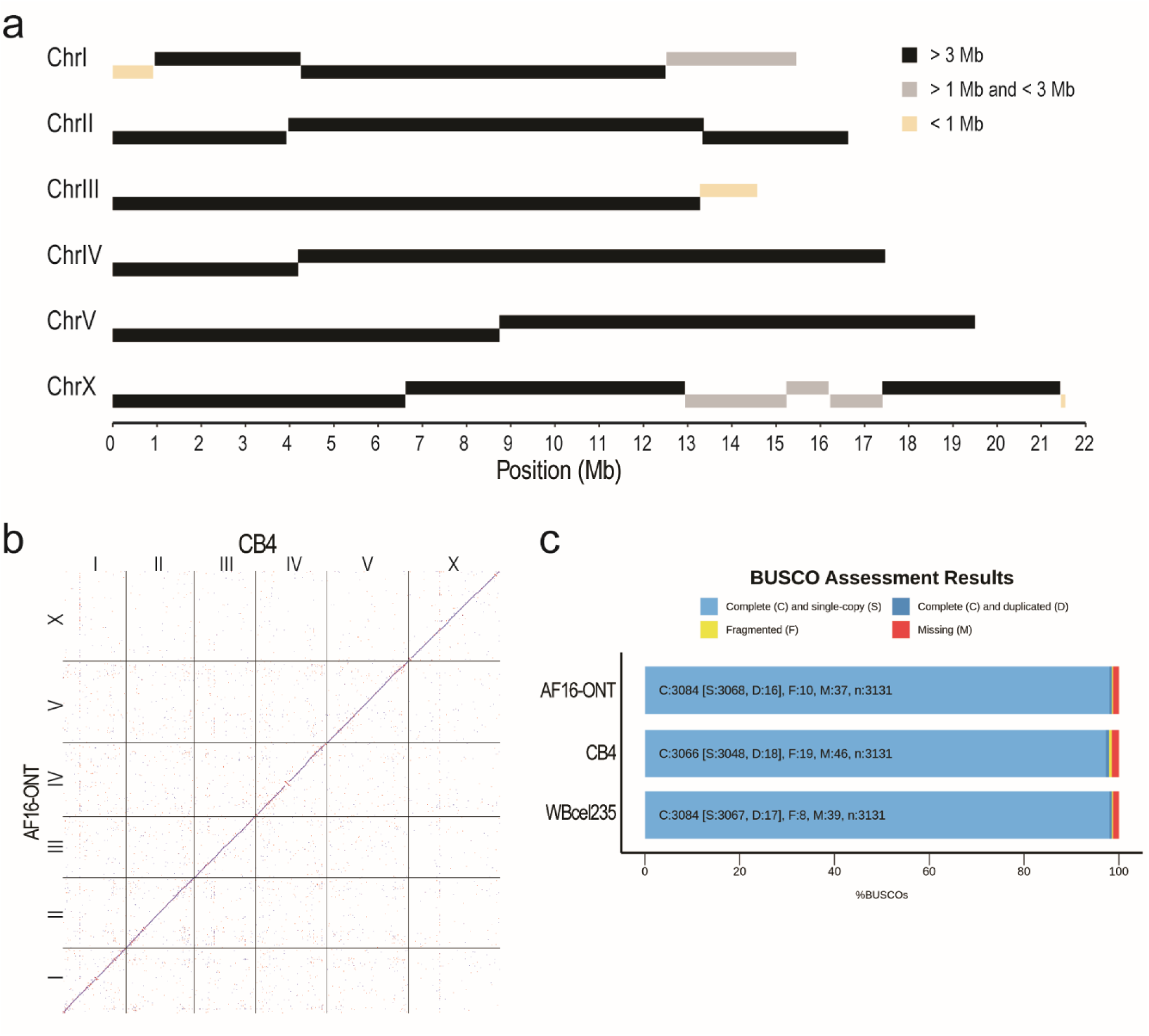
Evaluation of the ONT reads assembled *C. briggsae* AF16 genome. (a) Schematic representation of AF16 long reads assembled contig lengths. (b) Dot plot of corresponding chromosomes between CB4 and ONT reads assembled genome. (c) Bar chart with summary assessment for the proportion of genes present in three assembled genomes. AF16-ONT: the assembled *C. briggsae* draft genome in this study, WBcel235: the *C. elegans* N2 reference genome, CB4: the *C. briggsae* AF16 reference genome.

**Fig. 5.**
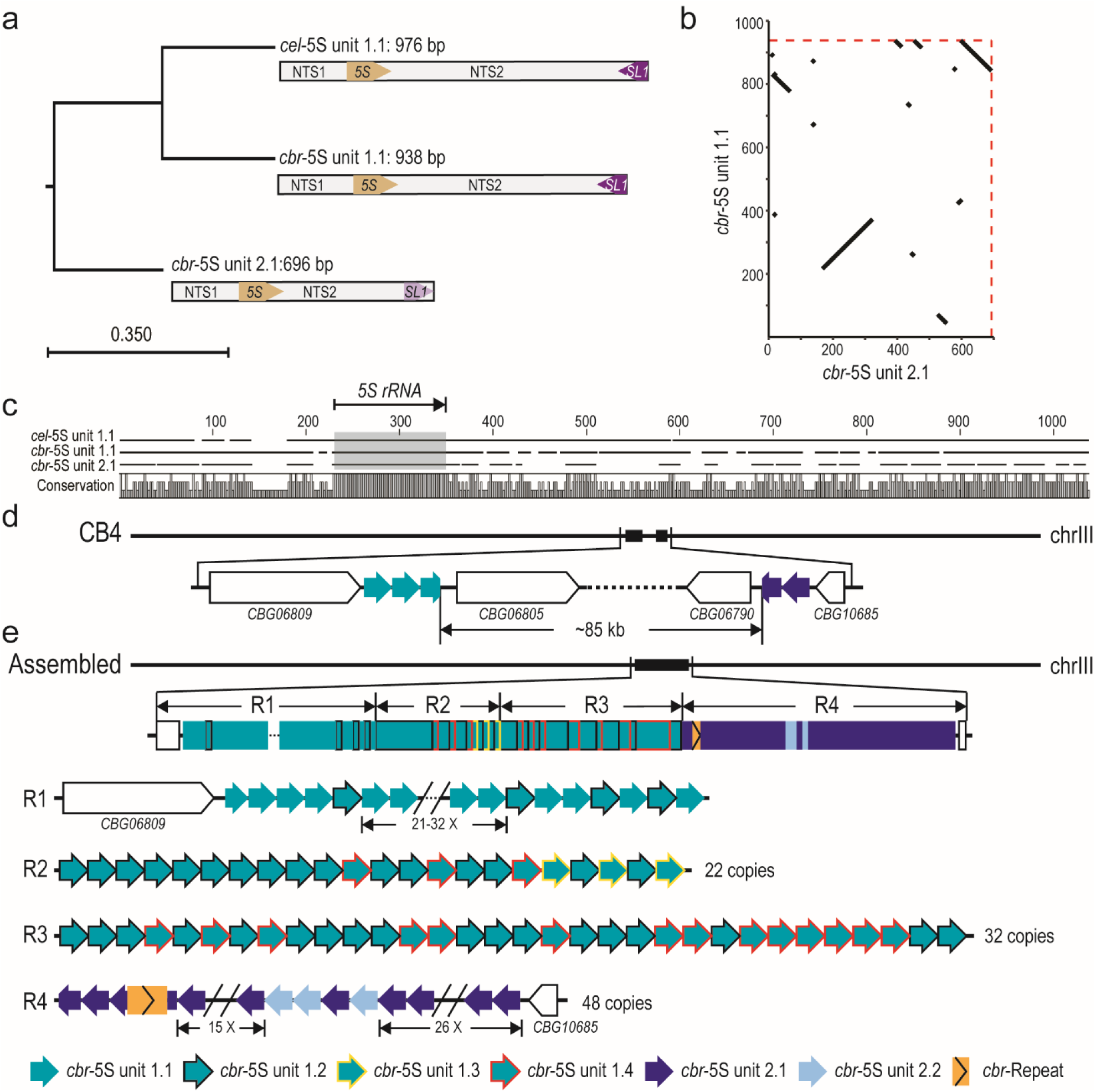
Characterization of the 5S rDNA units in *C. briggsae* AF16. (a) Phylogenetic tree of two divergent 5S rDNA units in *C. briggsae* (*cbr*) and the canonical *C. elegans* (*cel*) 5S rDNA unit. (b) Dot plot showing the sequence alignment between two *C. briggsae* 5S rDNA units. (c) Multiple sequence alignment of 5S rDNA units from *C. elegans* and *C. briggsae*. Alignments for the 5S rRNA gene is shaded in the grey box (indicated at the top). (d) A contig was misassembled into the rDNA cluster on chromosome III in the reference genome CB4. (e) Schematic representation of *C. briggsae* AF16 5S rDNA cluster annotated by ONT reads.

### Transposition of chromosome I end associated with 45S (18S-5.8S-26S) rDNA cluster in *C. elegans* genome

Unlike in yeast, the locus of 5S rDNA cluster is separated from that of the 18S-5.8S-26/28S rDNA cluster in nematode [2, 17], fly [42], mouse and human [43]. The 45S rDNA unit consists of an 18S, a 5.8S and a 26S rRNA gene interrupted by two internal transcribed <u>s</u>pacers (ITS1 and ITS2) in both *C. elegans* and *C. briggsae*. The *C. briggsae* 45S rDNA unit is roughly 300 bp longer than that of *C. elegans*, which was mainly contributed by the <u>e</u>xternal transcribed <u>s</u>equence (ETS) (Fig. 6a-c). The *C. elegans* 45S rDNA cluster is located at the right end of chromosome I. The ONT reads from all *C. elegans* N2-derived strains confirmed that the sequence between the 45S rDNA cluster and the telomeric sequences is partial ETS (Fig. 6d). Based the NGS reads of N2 genomic DNAs [35], most of the called variants using ONT reads (Fig. S5 and Table S6) resulted from INDELs in the homopolymer regions, in which ONT read sequence were known to be less accurate than in other regions, leading to our attempt to identify possible sequence variation within the cluster was not successful. In addition, all our ONT reads carrying either the left or right flanking sequences contain only partial 45S rDNA unit. This was mostly due to the relatively large size of the unit (∼7.2 kb in *C. elegans* and ∼7.5 kb in *C. briggsae*) and a relatively shorter 45S rDNA sequence-containing reads compared to other genomic positions (Fig. S6). Therefore, we were unable to identify any unique sequence in the cluster as an anchor to extend ONT reads deeper into the cluster from both boundaries. Although we are not certain whether there were any structural variations within the *C. elegans* 45S rDNA cluster, these ONT reads can be used to correct the boundary sequences of 45S rDNA cluster in *C. elegans* N2 and CB4856 strains (Fig. 6d). We observed a dramatic rearrangement event in the right boundary of CB4856 chromosome I relative to that of N2. For example, we identified an apparent transposition of the left end of chromosome IV to the right end of chromosome I (Fig. S7), which is consistent with a previous finding [32]. The transposed fragment underwent an uninterrupted duplication and transposition to the left end of the chromosome I along with its flanking rDNA sequences. A tandem array consisting of positioning sequence on X (pSX1) [44] was also found adjacent to the transposition site, but its origin was unclear.

**Fig. 6.**
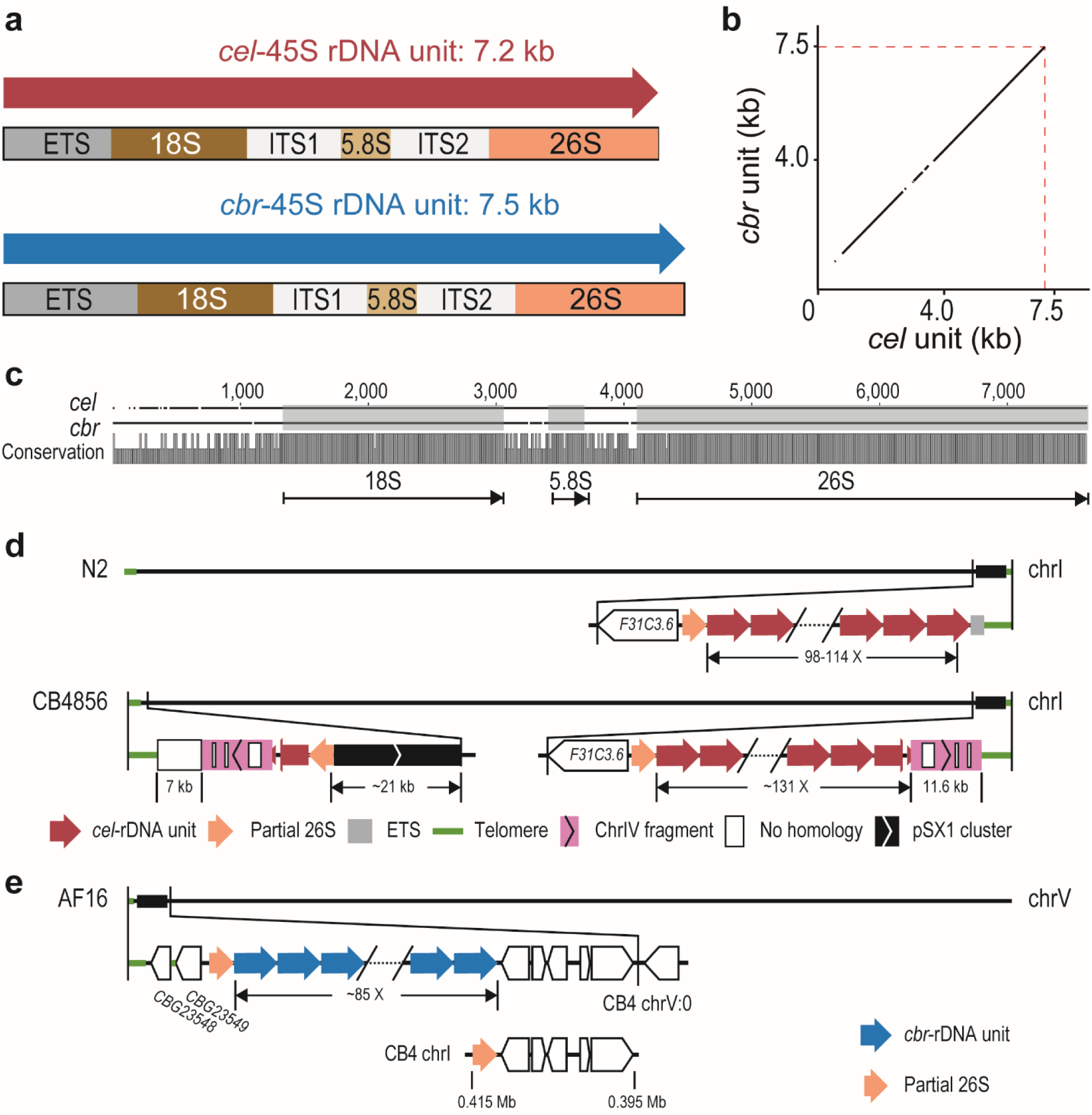
Comparison of 45S rDNA units and cluster between strains and species. (a) Comparison of 45S rDNA units between *C. elegans* and *C. briggsae*. (b) Dot plot showing the alignment of the 45S rDNA unit sequences between two species. (c) Pairwise sequence alignment of the 45S rDNA unit between two species. The 18S, 5.8S, and 26S RNA gene regions are shaded in grey. Conservation scores are shown at the bottom. (d) Schematics of the 45S rDNA cluster of *C. elegans* N2 and CB4856 annotated by ONT reads. In N2, the cluster left and right boundaries are flanked by partial 26S rRNA sequences and a partial ETS, respectively. In the 45S rDNA-containing region in *C. elegans* CB4856, the 45S rDNA cluster is located at the right end of chromosome I while fragmented 45S rDNA sequences along with other sequences are located at the left end. The estimated copy number of the unit is shown. Note that both the chromosome left and right ends are flanked by a ∼11.6 kb fragment derived from the left end of chromosome IV (pink, see Fig. S7), which is interrupted by some no homologous sequences (white box). A pSX1 cluster is also found adjacent to 45S rDNA. (e) Schematics of the *C. briggsae* AF16 genomic regions containing the 45S rDNA annotated by ONT reads in this study. Reconstructed 45S rDNA cluster is located at the left end of chromosome V containing about 85 copies of 45 rDNA unit. Bottom: A misassembled contig containing partial 26S rRNA gene sequences and 5 protein coding genes was assigned to chromosome I in CB4.

In the *C. briggsae* genome assembly CB4 [24], the 45S rDNA-containing sequences were fragmented in various contigs with unknown chromosome linkage (Fig. 6e and Fig. S2d). The Hi-C data [28] and our ONT reads supported a single location of the 45S rDNA cluster at the left end of the chromosome V (Fig. 6e and Fig. S4b). We further evaluated the validity of the estimated copy number of 45S rDNA unit by mapping of our ONT reads against the 45S rDNA cluster consensus sequences incorporated into our newly generated *C. briggsae* genome. The changes in reads coverage were consistent with the estimation of 45S rDNA copy number (Fig. S2d).

### The genomic environment of rDNA cluster is compatible with RNA Pol II transcriptionally

Eukaryotic cells use at least three RNA polymerases, i.e., RNA polymerase I (Pol I), Pol II, and Pol III, which produce 18S/5.8S/26(28)S rRNA, mRNA, and 5S rRNA, respectively. Given that all the rDNAs transcribed by the RNA Pol I and III are localized at two distinct loci consisting of rDNA and some other repetitive sequences only but depleted of any protein-coding sequences in both *C. elegans* and *C. briggsae* genomes, and yeast mutant lacking rDNA locus can be rescued by forced expression of rRNAs by RNA polymerase [45], we wondered whether the two rDNA clusters are permissive to RNA Pol II transcriptionally. To this end, we generated multiple transgenic lines carrying a single copy of insertion within or outside the rDNA cluster expressing a fluorescence marker using *unc-119* mutant [37]. In the transgenic animals, a complete rescue of uncoordinated (Unc) phenotype along with an apparent expression of the reporter in various parts of the soma indicates the native rDNA cluster regions are transcriptionally compatible with Pol II in the somatic tissues (Fig. S8). However, despite the expression of the reporter in soma, germline, and early embryo when it was inserted outside of the rDNA cluster, the expression in germline and early embryo was absent for the same reporter inserted within the rDNA cluster, suggesting that the genomic environment of rDNA cluster may not be accommodative to the expression in germline and early embryo.

## Discussion

Rapid development in sequencing technologies that can produce ultra-long reads makes it possible for resolving the structures of complex genome regions, including those consisting of tandem repetitive sequences. These sequences represent the “dark matter” of the existing genomes, including human genome [46]. One of the key advantages of the long reads is their ability to span repetitive sequences, allowing *de novo* assembling of the repetitive region or scaffolding of the existing contigs generated from NGS reads. Aided by the long reads, resolving the structure of highly repetitive regions, including rDNA cluster, centromere, telomere, or chromosomal rearrangement, becomes within reach. Our analyses of rDNA cluster structures using ONT long reads in both *C. elegans* and *C. briggsae* provide insights into the intra- or inter-species dynamics of rDNA clusters, which demonstrate an unusually high rate of structural and sequence variations inside the 5S rDNA cluster in the *C. elegans* N2 strain compared with its distantly related *C. elegans* CB4856 strain and the *C. briggsae* AF16 strain. The results suggest that *C. elegans* N2 strain may be at a disadvantage in maintaining the structure and stability of its rDNA cluster relative to other strains or *Caenorhabditis* species. This may have complications in its fitness, which warrants further investigation.

### The power of ONT read in resolving tandem repeats

Repetitive sequences especially those tandem repetitive ones are problematic for genome assembly. The *C. elegans* genome has been claimed as a “finished” genome with no gap due to its homozygosity and relatively small size [21]. However, the annotation of its genomic regions involving rDNA sequences is far from completion. For example, except for the boundary sequences, the previous sequencing methods failed to establish the genomic arrangement of the rDNA units and their variations [21,30,31]. Meanwhile, the existing *C. briggsae* genome assembly is far more fragmented than the *C. elegans* one. Despite various efforts have been made to improve the genome assembly of *C. briggsae* [19,23,24,47,48], none of them has been able to reliably resolve the structure and genomic localization of rDNA cluster. Aided by the ONT reads of high coverage, the genomic localization was readily resolved for both 5S rDNA and 45S rDNA clusters in *C. briggsae* (Fig. 5e and Fig. 6e). Our method of using Nanopore sequencing in resolving complex genomic structures and repetitive regions is readily applicable to rDNA cluster in other species. Consistent with this, taking advantage of ONT reads, the entire human X and Y chromosomes were assembled from telomere to telomere using genomic DNAs of an isogenic cell line [49, 50].

Most nematode genomes were assembled as contigs using shotgun sequencing method with NGS reads [51], which is also the case for many other species, leading to the absence of genomic parts consisting of tandem repetitive sequences. Given the decreasing sequence costs of ONT reads, it is feasible to re-sequence or improve numerous existing genomes especially for those of human and model organisms as well as economically significant species using the reads produced by ONT or other sequencing platforms such as PacBio High-Fidelity (HiFi). Given a relatively lower read accuracy of ONT reads than NGS reads, it would be ideal to simultaneously generate new or use the existing NGS reads to correct the nucleotides of a *de novo* genome assembly generated with ONT reads only. This would give rise to a highly accurate genome in terms of nucleotide and chromosome continuity. Given that most of the existing genomes were generated using NGS reads, leading to a fragmented assembly because of the presence of highly repetitive sequences, the TGS is expected to play a significant role in genome finishing or improvement in the years to come.

### Failure of recovering any ONT read that spans the entire 5S rDNA cluster suggests its complex structure

Given the ONT read length up to 196 Kbps (Table 1), the estimated copy number (Table S7), and relatively small size of the 5S rDNA unit, we reasoned that there were at least some ONT reads that were able to span the entire region from the left boundary of the 5S rDNA cluster to the “anchoring” sequence, i.e., the unique variants of 5S rDNA unit or the transgenes landed inside the cluster (Fig. 2a, b). However, we failed to recover any such ONT reads, suggesting that there could be some complex structural barriers that prevented the extension of DNA strand during Nanopore sequencing. Consistent with this, we observed a relatively smaller average read length of ONT reads associated with rDNAs than those independent of rDNAs (Fig. S6). For example, in the strain ZZY0603, which carries a transgene inside the 5S rDNA cluster (Fig. 2b), the ONT reads associated with the transgene contained up to 52 copies of canonical 5S rDNA unit on the left side of the transgene. However, no read was found to span the entire region from the left boundary of rDNA cluster to the transgene. This was unexpected because the entire unresolved part within the R1 region was estimated to carry a total of 34-60 copies of 5S rDNA unit with 31 copies located on the right side of transgene (Fig. 2b). Similarly, in the ONT reads of ZZY0600, which carried a transgene next to the sixth copy of the 5S rDNA unit away from the left boundary of the 5S rDNA cluster, the ONT reads associated with the transgene carried a maximum of 44 copies of 5S rDNA unit on the right of the transgene (Fig. 2b). Again, no read was found to span the entire region from the anchoring 5S rDNA variant (unit 1.6) to the transgene. Hence, we postulate that rDNAs in this region may undergo active DNA replication or amplification, which prevented sampling of longer DNA fragment for sequencing. For example, at replication fork, the rDNAs undergoing active replication are single-stranded [8], which would be vulnerable to DNA shearing during DNA extraction. Consequently, the ongoing replication in rDNA may hinder the extension of the ONT reads, which led to the absence of the long reads spanning the entire active region. Alternatively, the failure of ONT read to span the entire region was likely caused by a complex tertiary structure of the highly repetitive DNA sequences, which might be difficult to be opened up by the helicase during Nanopore sequencing and blocked the sequencing pore, leading to early termination of ONT sequencing process.

### Uncoupled 5S rDNA and 45S rDNA copy number between developmental stages at organism level

The copy numbers among the 5S, 5.8S, and 28S rRNA genes, which encode rRNAs that constitute the ribosomal large subunit, were thought to be highly correlated [52, 53]. Given the differential transcriptional efficiencies between cell types and the storage of 5S rRNA in ribosome-free particles [54], the copy numbers of rRNA genes may not necessarily show concerted change at organism level although they could be coupled in a particular cell type. For example, the estimated copy numbers between 5S rDNA and 45S rDNA appeared to be uncoupled (Table S8). Copy number of the 5S rDNA unit increased from 116, 169, to 184 from EMB, L1, and YA stage, whereas the copy number of 45S rDNA unit reached the highest level at L1 stage (114 copy) compared with 98 and 103 copies at EMB and YA stages, respectively. Although this result is consistent with a previous finding with mutated *C. elegans* NGS data [5], it is inconsistent with the data from human and mouse [52], suggesting differential regulations of the overall dosage of 5S and 45S rDNAs between nematodes and mammals.

Availability of ultra-long reads from ONT or PacBio platforms is expected to accelerate the generation of complete genome sequence from telomere to telomere. With these technologies, the human chromosome 8, X, and Y were assembled with no gap albeit with some manual corrections [55–57]. With ultra-long reads of a higher read accuracy, the structure of 45S rDNA and other highly repetitive regions such as centromeres and telomeres are expected to be resolved in the coming years, leading to a gap-free genome, in human, model organisms and economically significant species in the years to come.

## Methods

### Sequencing library preparation and ONT sequencing

For *C. elegans* wild isolates, genomic DNAs were extracted from the mix-staged embryos (EMB), early-stage larvae (L1) and young adults (YA) of N2 strain (shipped from Waterston laboratory, Seattle, WA, USA in 2010) or from the mix-staged animals of CB4856 strain. For *C. elegans* transgenic strains, genomic DNAs were extracted from the homozygous mix-staged animals with the following genotypes: ZZY0600 (*unc-119*(*tm4063*) III; *Is*[*sel-8p*::HIS-24::GFP::*pie-1* 3’ UTR, *unc-119*(+)] V), ZZY0603 (*unc-119*(*tm4063*) III; *Is*[*dsl-1p*::HIS-24::GFP::*pie-1* 3’ UTR, *unc-119*(+)] V), and ZZY0653 (*unc-119*(*tm4063*) III; *Is*[*his-72p*::mCherry::HIS-24::*pie-1* 3’ UTR, *unc-119*(+)] I), each carrying a single-copy of transgene in rDNA cluster. For *C. briggsae* wild isolate, genomic DNAs were extracted from AF16 young adults. Animal synchronization was performed as described [58]. Before harvesting, the *C. elegans* and *C. briggsae* animals were maintained on plates of 1.5% <u>n</u>ematode <u>g</u>rowth <u>m</u>edium (NGM) seeded with *E. coli* OP50 at room temperature and in a 25°C incubator, respectively. Genomic DNAs were extracted from animals with PureLink Genomic DNA Mini Kit (Invitrogen) using siliconized tubes and pipette tips to minimize shearing. 4 µg purified DNAs from each sample were used for library preparation using Genomic DNA by Ligation Kits SQK-LSK108 (Oxford Nanopore Technologies) for N2 and ZZY0653, and Ligation Kits SQK-LSK109 (Oxford Nanopore Technologies) for the remaining strains. Sequencing was performed on GridION X5 or MinION with R9.4.1 flow cell (FLO-106, Oxford Nanopore Technologies) using default parameters.

### Sequence acquisition and alignment

Base-callings were performed using Guppy (v3.1.5, Oxford Nanopore Technologies) using the high-accuracy configuration (HAC) model. All the base-called reads from each library were pooled for analysis of read length distribution with SeqKit (v0.10.2) [59]. The reads were aligned against the *C. elegans* N2 genome assembly (WormBase WBcel235) [34] or the *C. briggsae* AF16 genome assembly (CB4) [24] with Minimap2 (v2.17) [60] using default parameters for ONT reads. Read average coverage was calculated from the BAM file using SAMtools depth [61]. The ONT reads of *C. elegans* VC2010, a wild-type strain derived from N2, were downloaded from European Nucleotide Archive (ENA) with accession numbers PRJEB22098 [31]. The ONT and PacBio reads from *C. elegans* strain PD1074, a wild type strain derived from VC2010, were downloaded from Sequence Read Archive (SRA) database with accession number SRR7594463 and SRR7594465, respectively [30]. The ONT reads of VC2010 and PD1074 were used for identifying lab-specific variations in the rDNA unit and its genomic organization. The PacBio reads of *C. elegans* CB4856 were downloaded from the SRA database with accession number SRR8599837 [32].

For short NGS reads of *C. elegans* N2 and CB4856, the alignment BAM files were downloaded from *Caenorhabditis elegans* Natural Diversity Resource (CeNDR) project [35]. Short NGS reads were aligned to WBcel235 using BWA (v0.7.17) [62] with default parameters. The *C. briggsae* SLR reads and Hi-C reads were downloaded from the SRA database with accession number SRR6384296 and SRR6384332, respectively [23, 48].

### Identification of variation in 5S rDNA units

*C. elehans* ONT reads with rDNA sequences were aligned against one copy of *cel*-5S unit 1.1 with Minimap2. From the CIGAR strings in the generated SAM file, to minimize the INDELs resulted from base-calling errors from homopolymers and simple repeats, only the INDELs longer than 3 bp were kept for copy counting with custom scripts. After normalization with genome-wide read coverage, the normalized INDEL count higher than one copy was considered as a potential new INDEL variant. Using the strain-specific BAM files generated with NGS read alignment against the N2 reference genome produced previously [35], *C. elegans* N2 and CB4856 NGS reads mapped to the 5S rDNA region were separately extracted and then individually mapped to a single *cel*-5S unit 1.1 in the same way as that for the ONT reads. SNP calling was performed with BCFtools [63] using the NGS reads stated above. The presence of *cel*-5S unit 2 was established by a 4-bp deletion relative to the *cel*-5S unit 1.1. The *cel*-5S unit 3 was established by the presence of a 30-bp deletion in the NGS and ONT reads relative to the *cel*-5S unit 1.1 in N2 strain. This deletion was absent in the NGS and ONT reads of CB4856.

To investigate whether the 30-bp deletion in the *cel*-5S unit 3 are present in all *C. elegans* wild-isolates, the NGS reads derived from 330 whole-genome shotgun libraries [35] were mapped against the sequences of *cel*-5S unit 1 and 3 using BWA. The reads that were uniquely mapped to the deletion junction for at least 12 bps at both flanking sides were extracted with SAMtools with parameters -q 30 -F 4. A strain was defined as *cel*-5S unit 3-containing if over 1% of total reads carried the deletion regardless of the total number of supporting reads, or if over 0.1% of total reads carried the deletion but with at least 10 supporting reads. The phylogenetic tree of the 330 strains produced previously [35] was visualized in R with ggplot2 and ggtree packages [64–66]. The variants of *C. briggsae* 5S rDNA units were identified similarly as in *C. elegans*.

### Reconstruction of rDNA clusters

Reconstruction of the *C. elegans* 5S rDNA cluster started with identifying all the ONT reads carrying the flanking sequences of the cluster, i.e. the *ZK218.23* as the left boundary, and the sequences from chrV: 17,133,740-17,137,381 (WBcel235) as the right boundary. These reads were iteratively extended into the cluster by performing SNP- and INDEL-based manual assembly through chromosome walking. Based on the pairwise alignment results using BLASTN [67], the consensus of rDNA cluster was generated using at least 10 supporting reads that contained the sequences of rDNA variants or other repeats as anchors from both DNA strands (Fig. S1). This step was reiterated till the exhaustion of all available ONT reads. To determine the potential structural variations among *C. elegans* N2-derived strains and between *C. elegans* strains, each 5S rDNA cluster was similarly assembled with strain-specific ONT reads. For assembly of 45S rDNA cluster in *C. elegans* N2, the right boundary was determined using the ONT reads containing both ETS and telomeric sequences (TTAGGC). For *C. elegans* CB4856 45S rDNA cluster, the right boundary was determined using the ONT reads containing telomeric sequences.

Reconstruction of the *C. briggsae* 5S rDNA cluster was started with two chromosome III contigs carrying a 5S rDNA sequence and genes next to rDNA sequences (*CBG06809* and *CBG10685*). The right boundary of 45S rDNA cluster was determined with the ONT reads carrying the rDNA sequence and those from its right boundary in CB4, which is located at the beginning of chromosome V. The 45S cluster left boundary was determined with the ONT reads carrying both 45S rDNA and telomere sequences.

### Draft genome assembly and quality assessment

To get a better reference genome for locating *C. briggsae* rDNA clusters, an AF16 draft genome was *de novo* assembled with ONT reads. Miniasm (v0.3) was run with AF16 young adult ONT reads. The generated contigs were polished with Racon (v1.4.10) [40] using two rounds of ONT reads and three rounds of SLR reads [23]. Bacterial genomes were manually excluded from the polished contigs. Remaining 21 contigs were scaffolded into chromosome level using CB4 as reference and interspaced with 1000 Ns. The final draft genome was aligned against CB4 using LAST (v1021) [68]. The completeness of the resulting AF16 draft genome assembly, CB4, and *C. elegans* N2 genome assembly WBcel235 was assessed in parallel using BUSCO (v4.0.2) [41] with nematoda_odb10 database.

### Estimation of rDNA copy number

For estimation of copy number of the *C. elegans* 5S rDNA units, ONT reads mapped to the region of the chrV: 17,110,000-17,430,000 (WBcel235) were extracted with SAMtools and were used for statistical analysis with SeqKit. The extracted reads were aligned against the 5S rRNA-coding sequence with BLASTN with option “-word_size 7”. Sequences with alignment length > 17 bps were kept for the downstream analysis. The copy number of the 5S rDNA units was estimated for each library by dividing the summed read lengths aligned to 5S rRNA by the product between 5S RNA gene length (119) and genome-wide read coverage. For estimation of copy number of the *C. elegans* 45S rDNA units, the reads mapped to the end sequence of the chromosome I (chrI: 15,057,500-15,072,434) were extracted and aligned against the ITS1. Sequences with alignment length > 21 bps were kept for further calculation. The 45S rDNA copy number in each library was estimated in a way similar to that for 5S rDNA.

For copy number estimation of *C. briggsae* 5S rDNA units, the reads mapped to the region of the chrIII: 10,555,000-10,660,000 (CB4) were extracted. The extracted reads were aligned against the 5S rRNA-coding sequences and two existing 5S rDNA units with BLASTN. Reads were retained for further analysis if their alignment sizes were bigger than 17, 170 and 170 bps for 5S RNA-coding sequence, *cbr*-5S unit 1.1 and *cbr*-5S unit 2.1, respectively. The copy number of 5S rDNA units was calculated in the same way as that in *C. elegans*. To extract all the reads mapped to the *C. briggsae* 45S rDNA cluster, a pseudo-chromosome was generated using chrI: 395,000-417,500 (CB4), which consisted of partial 26S rRNA coding sequence and its five protein-coding genes flanking the rDNA gene, and 100 copies of the *C. briggsae* 45S rDNA unit derived from SLR reads [23]. Reads mapped to the pseudo-chromosome were extracted and aligned against the *cbr*-ITS1 sequence with BLASTN. The estimated copy number of the 45S rDNA units was calculated in the same way as that in *C. elegans*.

### Validation of genomic localization and structure of assembled rDNA clusters

To validate the genomic localization of the assembled rDNA clusters in *C. elegans* N2 and *C. briggsae* AF16, the Hi-C sequencing data from L1 stage animals [28, 69] were employed to confirm the linkage between the rDNA clusters and their host chromosomes. For *C. elegans* reads, the rDNA pseudo-chromosome, which contains 50 copies of *cel*-5S unit 1.1 or 10 copies of *cel*-45S rDNA, was included into the reference genome for mapping of Hi-C reads. After trimming reads with Trimmomatic (v0.35) [70], the remaining reads were input to Juicer (v1.5) [71] with default parameters to find chromatin interactions between the rDNA pseudo-chromosome and host chromosomes. The density of interaction was normalized and visualized in R with circlize package (v0.4.7) [66, 72]. The linkage between the rDNA clusters and their host chromosomes in *C. briggsae* was performed in the same way as that in *C. elegans*. Specifically, the rDNA pseudo-chromosomes consisting of 50 copies of *cbr*-5S unit 1.1, or 50 copies of *cbr*-5S unit 2.1 and unit 2.2 with mixed arrangement, or 10 copies of *cbr*-45S rDNA, were individually included to the *C. briggsae* genome assembly CB4, respectively.

To evaluate the structure of the reconstructed 5S rDNA clusters, the existing rDNA cluster sequences in reference genome were replaced by the reconstructed rDNA sequence consisting of the minimum estimated copy number. The ONT reads were mapped against the modified reference genomes incorporated with the reconstructed rDNA sequence using Minimap2 with default parameters. The coverage within the reconstructed rDNA cluster regions was visualized in R with ggplot2 package [65].

### Molecular biology, transgenesis, and imaging

All promoter fragments were amplified from N2 genomic DNA with PCR primers listed in Table S9. The *miniMos* targeting vector pCFJ909 was modified to include a genomic coding region of *his-24* upstream of the GFP coding sequence to facilitate nuclear localization, as previously described [73]. The fragments were cloned into the modified pCFJ909 [36], resulting in a reporter cassette consisting of the sequences coding HIS-24::GFP or mCherry::HIS-24 and the sequence of *pie-1* 3’ UTR as described previously [74]. The vector was used for transgenesis with *miniMos* technique [36]. The transgene insertion site was determined using inverse PCR [36]. All micrographs were acquired with an inverted Leica SP5 confocal microscope equipped with two hybrid detectors at a constant ambient temperature of approximately 20°C.

## Data access

All base-called Nanopore reads from this study have been submitted to the NCBI BioProject database (https://www.ncbi.nlm.nih.gov/bioproject) under accession number PRJNA562392. The SRA and ENA accession number for each library is listed in Table S7. The sequences of rDNA units from this study were deposited in GenBank under accession number: MN519135 for *cel*-5S unit 1.1, MN519140 for *cel*-45S rDNA unit, MN519137 for *cbr*-5S unit 1.1, MN519138 for *cbr*-5S unit 2.1, and MN519141 for *cbr*-45S rDNA unit.

Variant sequences for each rDNA unit and custom scripts for analyzing INDELs and SNPs of ONT reads with rDNA sequences were deposited in GitHub https://github.com/qiutaoding/QD_nanoKit_py3/tree/master/rDNA.

## Acknowledgments

We thank Dr. Cindy Tan for logistic support and members of Zhao’s lab for helpful comments. Part of strains in this study were provided by the CGC, which is funded by NIH Office of Research Infrastructure Programs (P40 OD010440). The phylogenetic tree data for 330 *C. elegans* wild-isolates was kindly provided by Dr. Erik Andersen. This work was supported by General Research Funds (HKBU12100917, HKBU12123716, N_HKBU201/18, HKBU12100118) from Hong Kong Research Grant Council and HKBU Research Committee and Interdisciplinary Research Clusters Matching Scheme 2019/20 for 2017/18 to ZZ.

## Author Contributions

QD performed *C. elegans* CB4856, *C. briggsae* AF16, and transgenic animals Nanopore sequencing. QD and XR performed *C. elegans* N2 animal synchronization and Nanopore sequencing. LC and VWH generated the two transgenic strains. QD and RL performed primary data analysis. QD performed rDNA variants characterization and manual rDNA cluster assembly. RL performed Hi-C analysis. QD upload base-called data to SRA for data sharing. ZZ and RL coordinated the project. QD, RL, and ZZ drafted the manuscript.

---

**Fig. S1.**
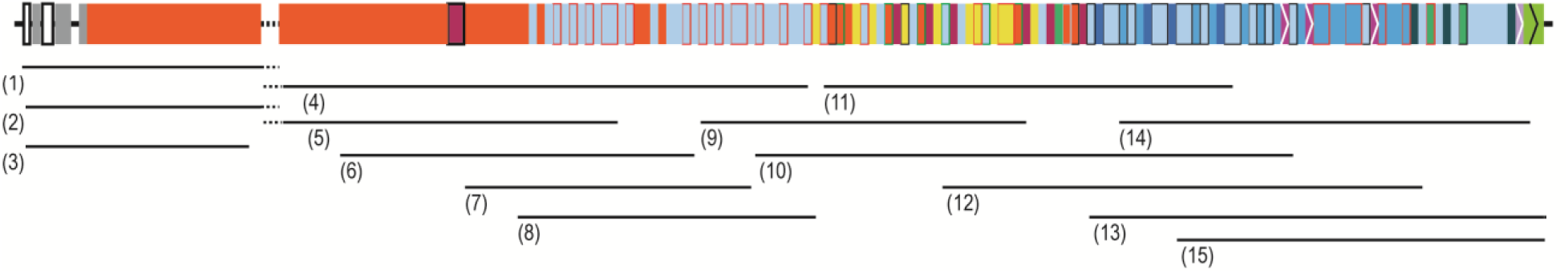
Illustration of tiling path method for reconstructing 5S rDNA cluster in *C. elegans* N2. The composition of 5S rDNA units are shown as bars color-coded as in Figure 1. The unsolved region (dash line) contains tandemly repeated *cel*-5S unit 1.1. Lines under the 5S rDNA cluster indicate the alignment of ONT reads. Read name and length are listed below. (1) 816e3c91-e978-404c-ad0c-0feac8208ddd, 63,658 bp. (2) 57556aff-f79a-4de4-81cc-d77ce3fec273, 48,846 bp. (3) 19aaf089-cf0c-4b24-9fae-2d14971bc320, 29,729 bp. (4) 28651bf0-8be1-4564-a740-ad17d0f260bd, 62,870 bp. (5) 77d84d3f-61ab-4fb1-9e4a-0fa291e6735d, 46,434 bp. (6) 594d2e66-ce5e-4b91-9f9e-7d6d24d9fbaa, 42,977 bp. (7) 7414a2d3-acdd-4c72-8509-e029ec1d7d13, 34,568 bp. (8) 367d1386-8328-4329-9ba6-c162993d8ede, 35, 021 bp. (9) 98b70228-6f3c-4e4b-ba71-b3631eea7acf, 38,624 bp. (10) 057d9497-6b73-4c82-9859-fd177a9930a2, 66,357 bp. (11) 9e96acd5-b022-4321-b922-bdbc7a33f7ab, 46,176 bp. (12) d42edfb5-7347-4c77-9d8c-23db3147a327, 50,719 bp. (13) 42938dc9-d10f-40cc-9558-cd6dade92882, 49, 434 bp. (14) c40285fc-c214-402d-b2c6-29b3d629c3f6, 52,513 bp. (15) bb18b8be-a27d-4119-aa91-b4b310f8cb28, 54, 031bp.

**Fig. S2.**
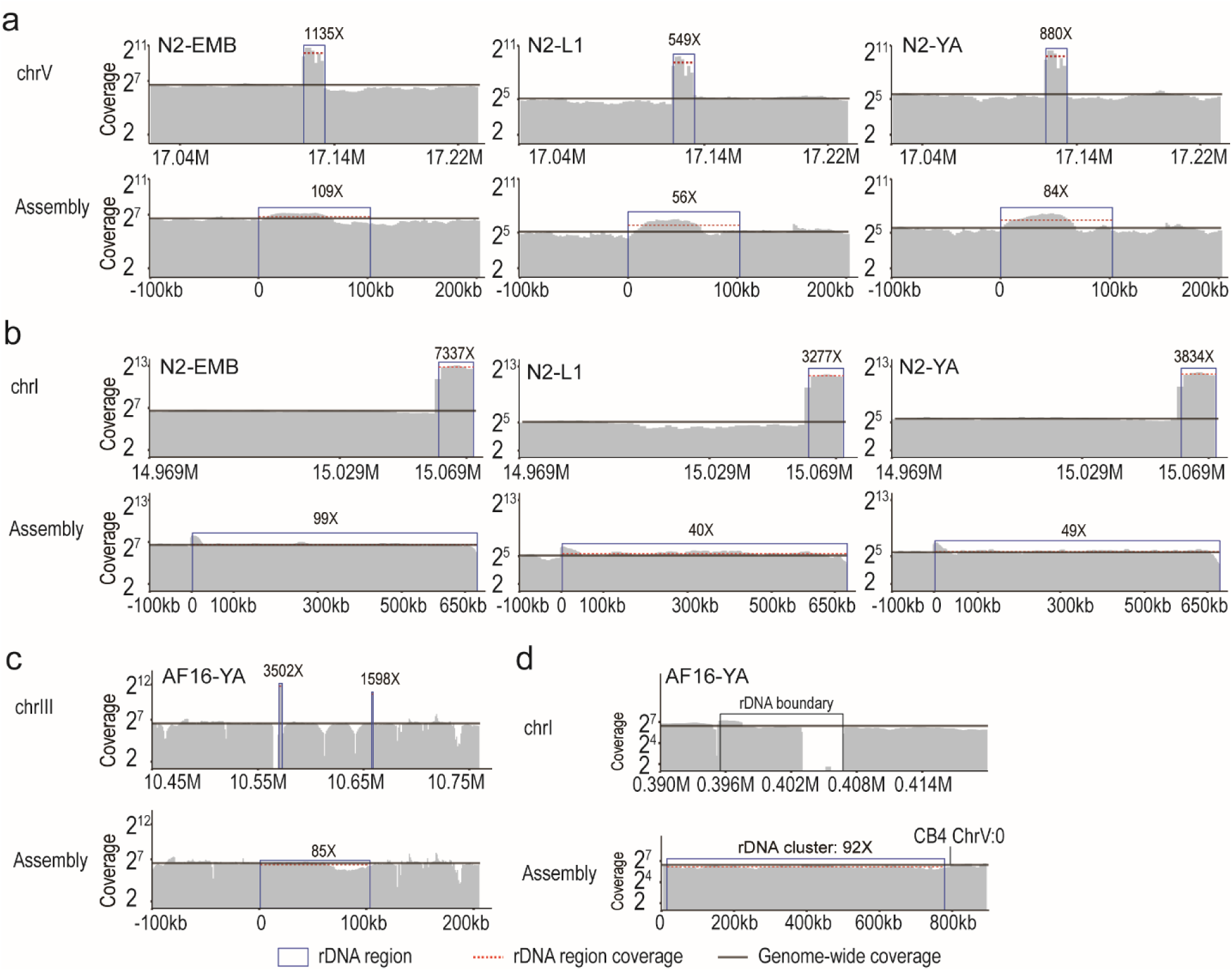
Coverage changes in rDNA clusters. (a) The coverage changes between the *C. elegans* N2 reference chromosome V 5S rDNA cluster region and ONT reads assembled 5S rDNA cluster consensus. (b) The coverage changes between reference chromosome I 45S rDNA cluster region and ONT reads assembled 45S rDNA cluster consensus. (c) The coverage changes between the *C. briggsae* AF16 CB4 chromosome III 5S rDNA cluster region and the ONT reads assembled 5S rDNA cluster consensus. (d) The coverage of misassembled 45S rDNA cluster flanking sequences on CB4 chromosome I and the ONT reads assembled 45S rDNA cluster-containing region on chromosome V.

**Fig. S3.**
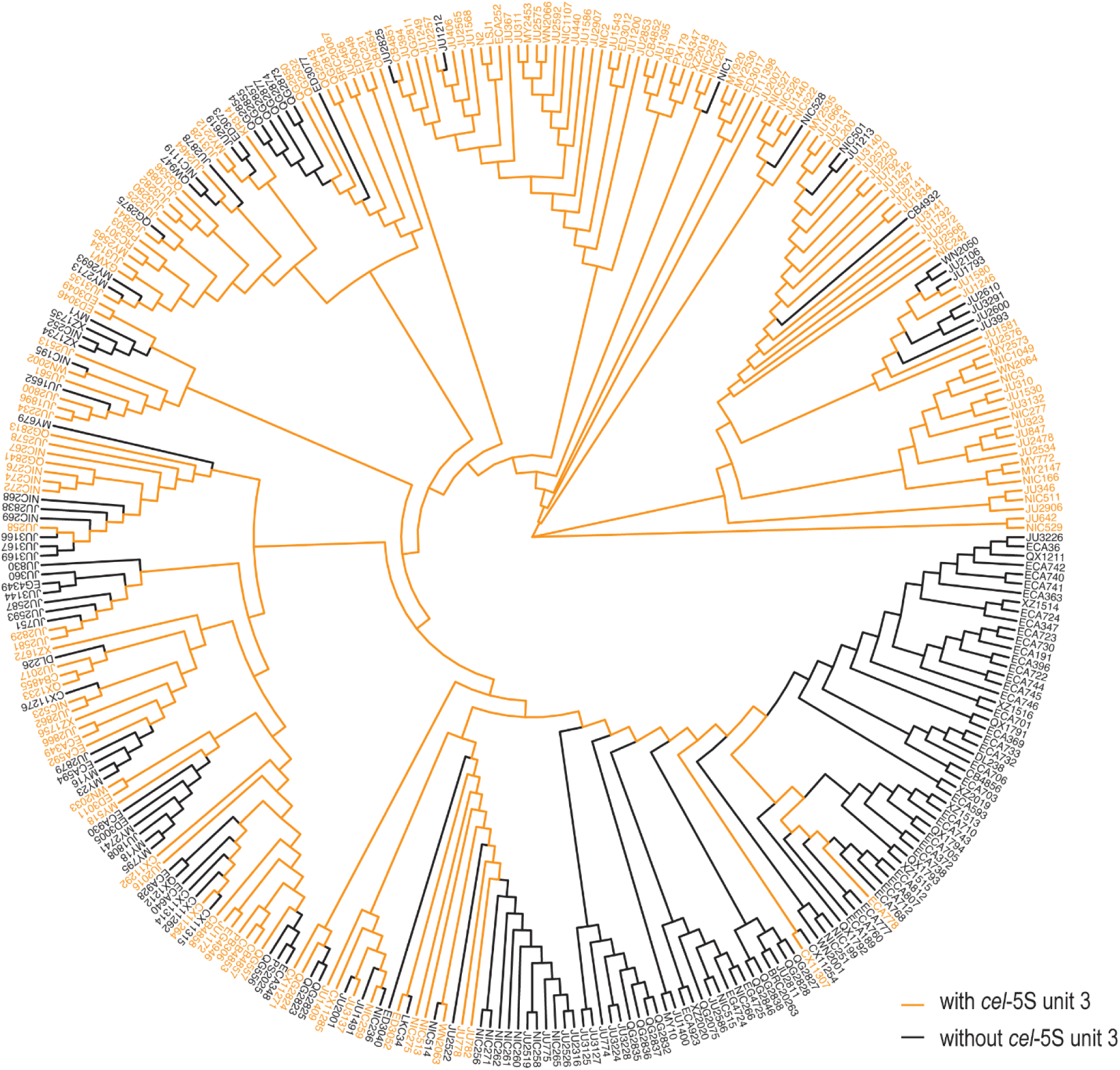
Presence of *cel*-5S unit 3 in the phylogenetic tree of 330 *C. elegans* wild-isolated strains. Branches marked with black color indicate that the *cel*-5S unit 3 is absent from NGS reads. See Table S5 for a detailed proportion of *cel*-5S unit 3 unique junctional reads in each strain.

**Fig. S4.**
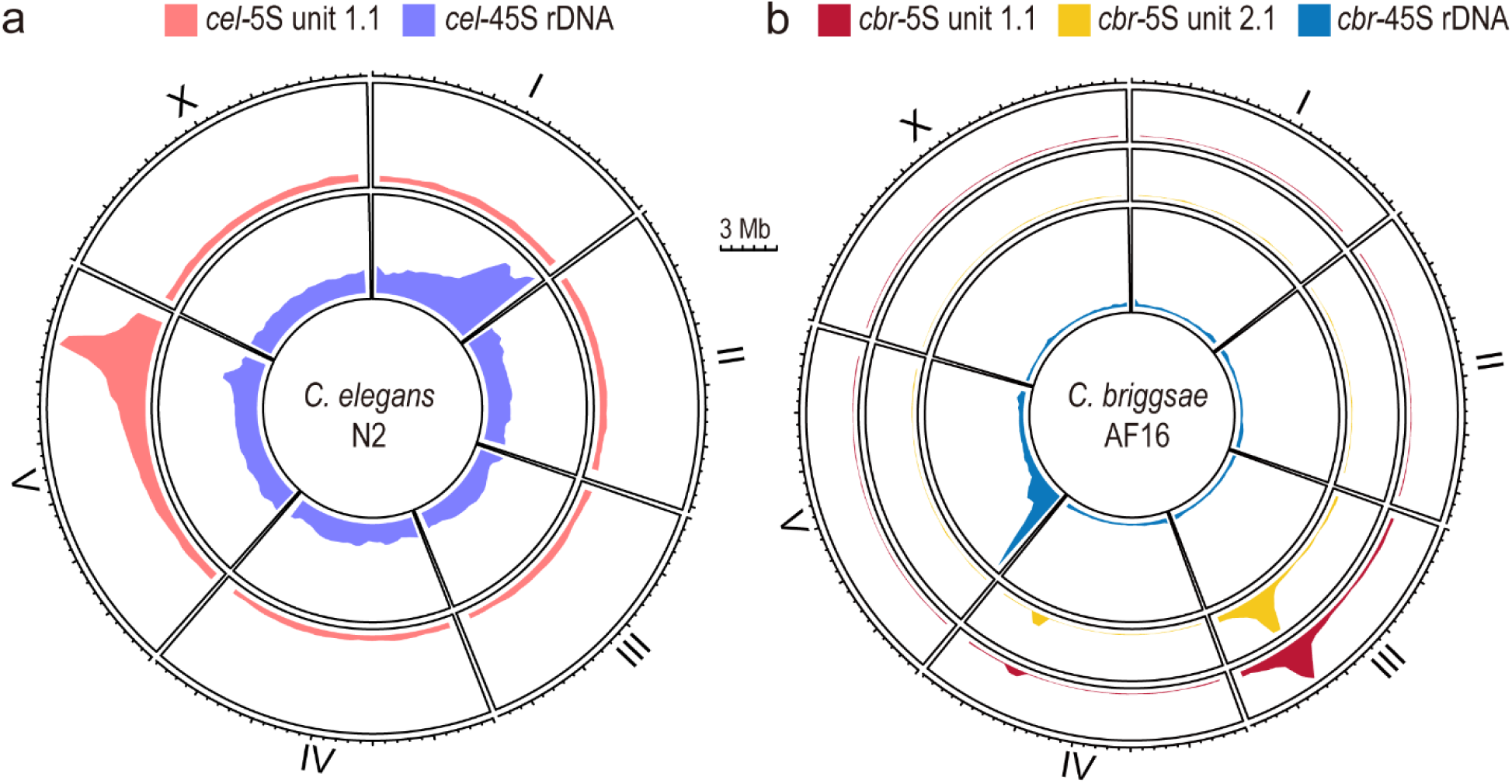
Hi-C interactions between rDNA clusters and chromosomes. (a) The *C. elegans* N2 genomic linkage density between chromosomes and the 5S rDNA cluster (pink in the outer circle) and 45S rDNA cluster (purple in the inner cycle). (b) The *C. briggsae* AF16 genomic linkage density between chromosomes and pseudo-chromosomes of *cbr*-5S unit 1 (red in the outmost cycle), *cbr*-5S unit 2 (yellow in the middle cycle) and 45S rDNA (blue in the inner cycle).

**Fig. S5.**
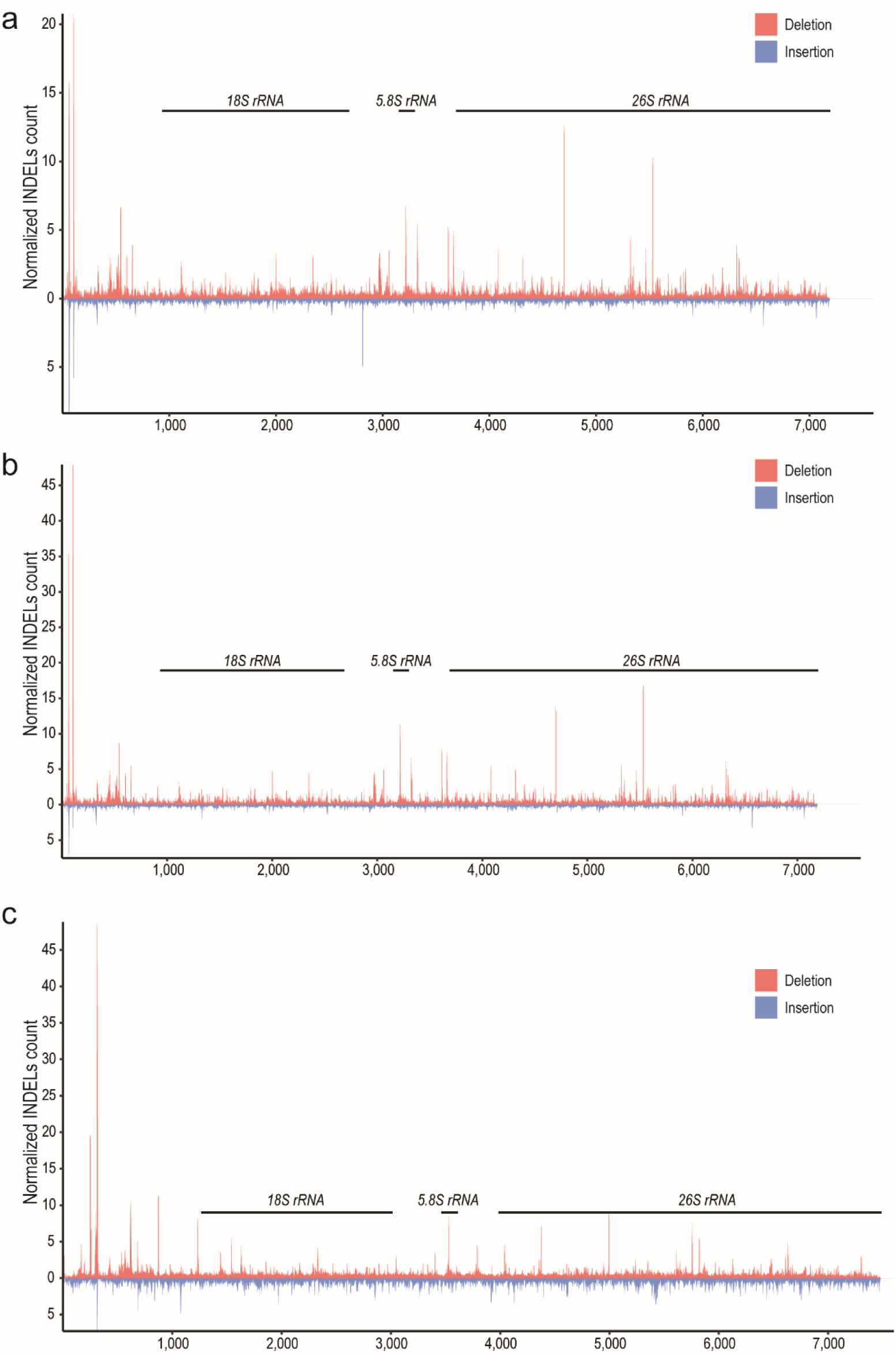
Large INDELs in 45S rDNA cluster of *C. elegans* and *C. briggsae*. Histogram plots show the normalized INDELs count of ONT reads mapped to a single copy of 45S rDNA sequences in *C. elegans* N2 (a) and CB4856 (b), and *C. briggsae* AF16 (c) 45S rDNA clusters.

**Fig. S6.**
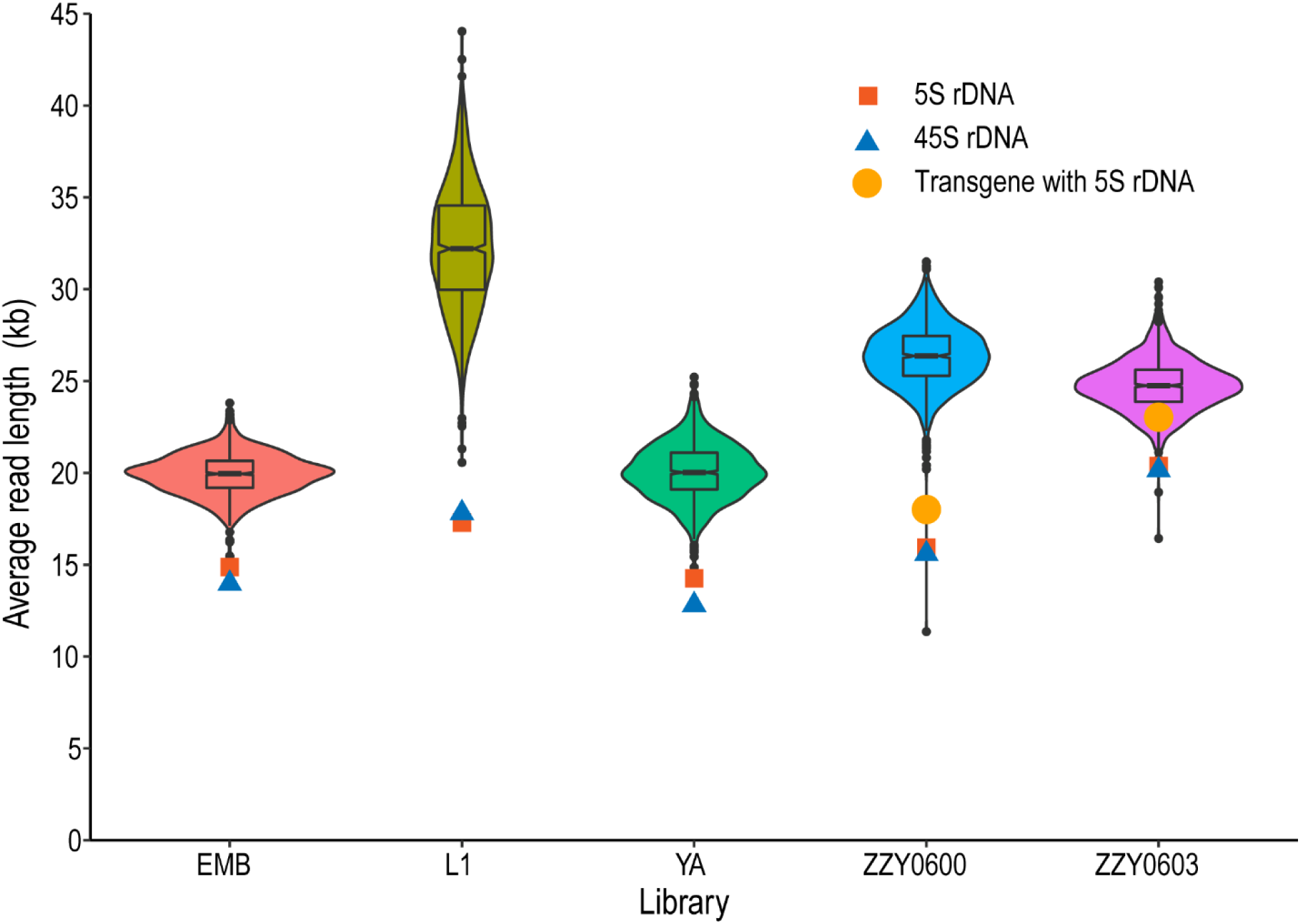
Smaller average read length in the rDNA clusters than in other genomic loci. Distribution of average read length from 999 positions in genome-wide positions and two rDNA clusters. Average lengths of ONT reads derived from 5S, 45S rDNA cluster, and transgene with rDNA sequences are denoted with a red square, blue triangle, and dark yellow dot, respectively. Box-plot shows the average read length ranging from Q1 quartile (25 percentiles) to Q3 quartile (75 percentiles).

**Fig. S7.**
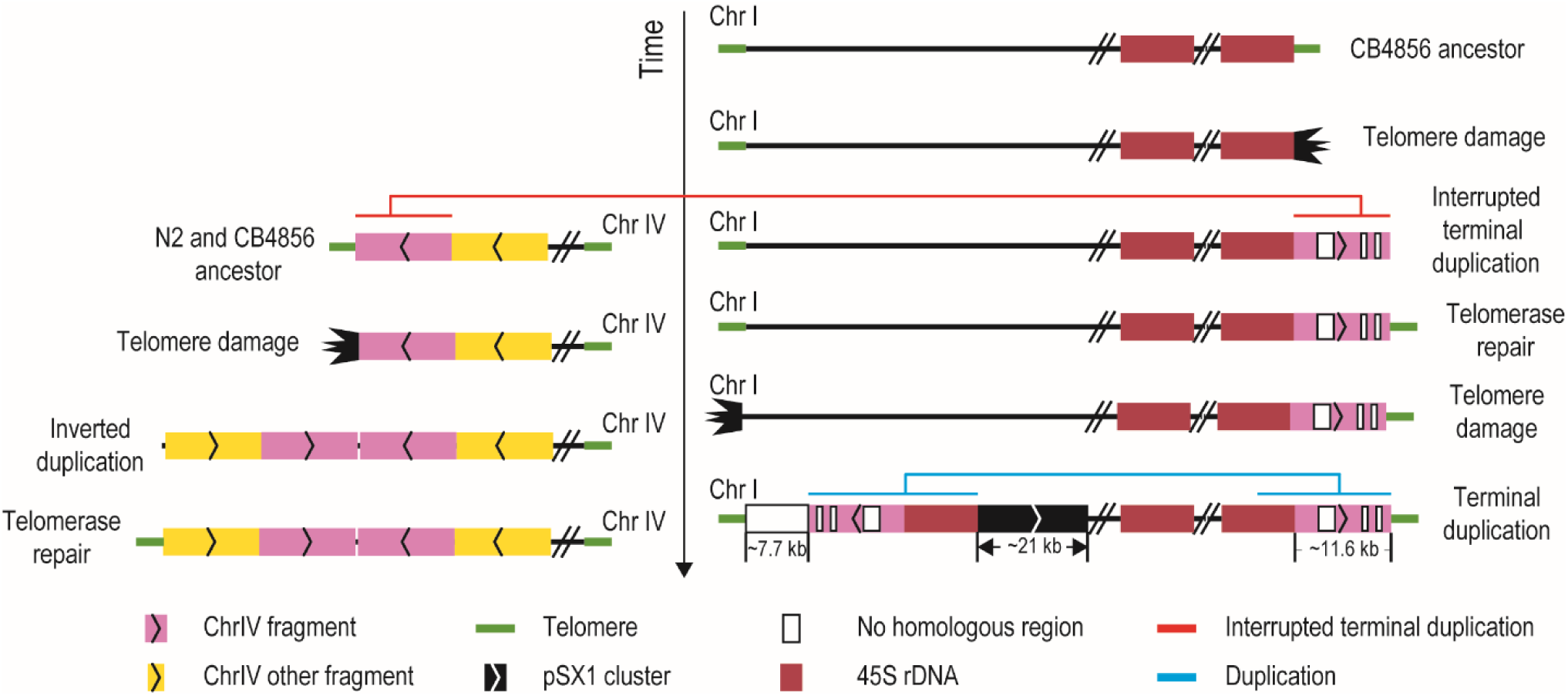
A terminal duplication model of chrIL, chrIR, and chrIVL ends in *C. elegans* CB4856 ancestor. The CB4856 ancestor underwent telomere damage at chrIR end and sequential telomere-damage repair by interrupted terminal duplication from Chr IVL subtelomere and telomere lengthening. Afterward, the ancestor chrIVL underwent a telomere crisis and repaired by an inverted duplication with telomerase repairing. Then, the ancestor met telomere damage at chrIL end and sequential damage repair by the new chrIR terminal duplication including partial 45S rDNA. The chrIL pSX1 cluster containing 124 copies of full-length pSX1 (172 bp) and 5 partial-length copies is ∼21 kb in length.

**Fig. S8.**
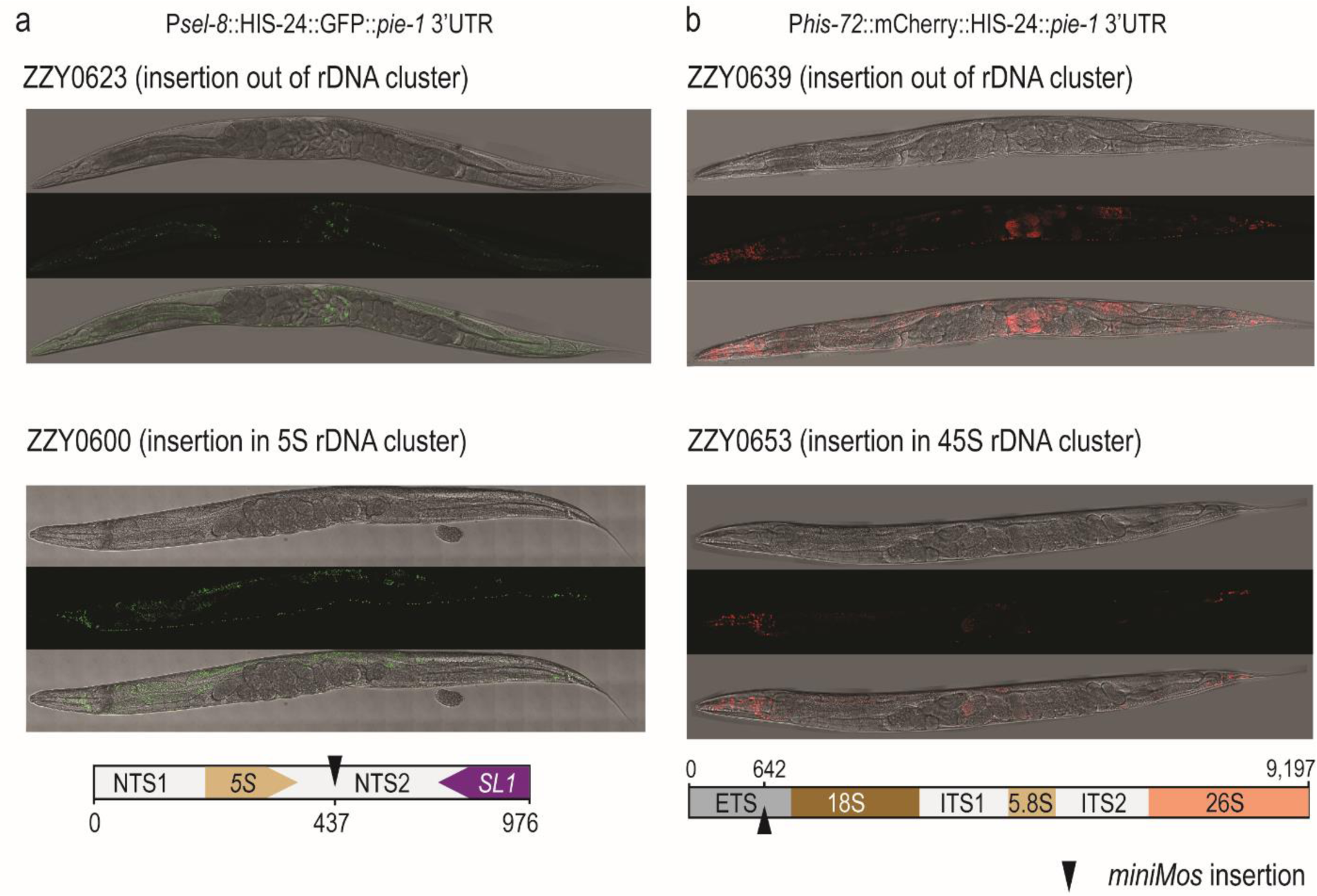
Altered expression pattern of transgenes in rDNA clusters. (a) Comparison of the expression of transgene inserted inside and outside of 5S rDNA cluster: Outside: GFP expression is found in embryos with 350 or more cells, and head and tail cells and some of the neurons in adult animals in ZZY0623. Inside: GFP expression is found in the head and tail cells and neuron nucleuses in ZZY0600 adult worm. (b) Comparison of the expression of transgene inserted inside and outside of 45S rDNA cluster: Outside, mCherry expression is found in late-stage embryos, mitotic germline, and ubiquitously in ZZY0639 adult animals; Inside, mCherry expression is found in late-stage embryos and head and tail cells in ZZY0653 adult animals. The transgene cassette in ZZY0653 was inserted into ETS in the opposite direction to the unit.

